# TAT-1, a phosphatidylserine flippase, affects molting and regulates membrane trafficking in the epidermis of *C. elegans*

**DOI:** 10.1101/2024.09.15.613099

**Authors:** Shae M. Milne, Philip T. Edeen, David S. Fay

## Abstract

Membrane trafficking is a conserved process required for the movement and distribution of proteins and other macromolecules within cells. The *Caenorhabditis elegans* NIMA-related kinases NEKL-2 (human NEK8/9) and NEKL-3 (human NEK6/7) are conserved regulators of membrane trafficking and are required for the completion of molting. We used a genetic approach to identify reduction-of-function mutations in *tat-1* that suppress *nekl*-associated molting defects. *tat-1* encodes the *C. elegans* ortholog of mammalian ATP8A1/2, a phosphatidylserine (PS) flippase that promotes the asymmetric distribution of PS to the cytosolic leaflet of lipid membrane bilayers. CHAT-1 (human CDC50), a conserved chaperone, was required for the correct localization of TAT-1, and *chat-1* inhibition strongly suppressed *nekl* defects. Using a PS sensor, we found that TAT-1 was required for the normal localization of PS at apical endosomes and that loss of TAT-1 led to aberrant endosomal morphologies. Consistent with this, TAT-1 localized to early endosomes and to recycling endosomes marked with RME-1, the *C. elegans* ortholog of the human EPS15 homology (EH) domain-containing protein, EHD1. TAT-1, PS biosynthesis, and the PS-binding protein RFIP-2 (human RAB11-FIP2) were all required for the normal localization of RME-1 to apical endosomes. Consistent with these proteins functioning together, inhibition of RFIP-2 or RME-1 led to the partial suppression of *nekl* molting defects, as did the inhibition of PS biosynthesis. Using the auxin-inducible degron system, we found that depletion of NEKL-2 or NEKL-3 led to defects in RME-1 localization and that a reduction in TAT-1 function partially restored RME-1 localization in NEKL-3–depleted cells.

**ARTICLE SUMMARY:** Endocytosis is an essential process required for the movement of proteins and lipids within cells. NEKL-2 and NEKL-3, two evolutionarily conserved proteins in the nematode *Caenorhabditis elegans*, are important regulators of endocytosis. In the current study, the authors describe a new functional link between the NEKLs and several proteins with known roles in endocytosis including TAT-1, a conserved enzyme that moves lipids between the bilayers of cellular membranes. As previous work implicated NEKLs in developmental defects and cancer, the present study can provide new insights into how the misregulation of endocytosis affects human health and disease.

## INTRODUCTION

Endocytosis is a highly conserved cellular process required for the uptake of extracellular molecules, the regulation of cell signaling, and interactions with the surrounding environment (DAUTRY-VARSAT AND LODISH 1984; MELLMAN 1996; DOHERTY AND MCMAHON 2009; GRANT AND DONALDSON 2009; SIGISMUND *et al*. 2012; BARBIERI *et al*. 2016; SIGISMUND *et al*. 2021). Moreover, by balancing exocytosis, endocytosis is essential for maintaining a consistent cell volume and plasma membrane surface area (BATTEY *et al*. 1999). During endocytosis, ligands, receptors, and other membrane-associated proteins, referred to as cargoes, are targeted to distinct membrane-bound compartments within the cell where they may be stored, degraded, or returned to the plasma membrane or organelles (MELLMAN 1996; KELLY AND OWEN 2011; CULLEN AND STEINBERG 2018).

Extracellular cargoes are internalized through several distinct mechanisms including clathrin-dependent and -independent forms of endocytosis (MCMAHON AND BOUCROT 2011; SANDVIG *et al*. 2018). Cargo uptake is typically followed by transport via vesicles to early endosomes (also referred to as sorting endosomes), which direct the flow of individual cargoes to specific locations (NASLAVSKY *et al*. 2003; JOVIC *et al*. 2010; NASLAVSKY AND CAPLAN 2018). Early endosomes may also undergo a process of maturation leading to the formation of late endosomes, where cargoes can be further sorted or retained for degradation within lysosomes.

After sorting at early endosomes, cargoes may be sent back to the plasma membrane through several distinct routes, collectively referred to as endocytic recycling pathways (CULLEN AND STEINBERG 2018; NASLAVSKY AND CAPLAN 2018). Most well understood is “slow” recycling, whereby cargoes are segregated within early endosomes and transported via vesicles to recycling endosomes (GRANT AND DONALDSON 2009; O’SULLIVAN AND LINDSAY 2020). There is also evidence that early endosomes can mature and take on the characteristics of the endocytic recycling compartment, although both processes likely co-occur (HUOTARI AND HELENIUS 2011; NASLAVSKY AND CAPLAN 2018). Once in the recycling compartment, cargoes are further segregated and ultimately exit via vesicles bound for fusion with specific plasma membrane domains or internal compartments (THOMPSON *et al*. 2007; HSU *et al*. 2012). How cells control the sorting and recycling of key cargoes is incompletely understood and can differ dramatically between cell types and under distinct physiological conditions (LOPEZ-HERNANDEZ *et al*. 2020). In addition, most studies on membrane trafficking have been conducted using dissociated cells in culture, which may differ with respect to endogenous processes that are carried out within tissues and intact organisms (GRANT AND SATO 2006; SCHMID *et al*. 2014).

Membrane lipid composition—along with regulators of membrane lipids—has recently been recognized to play major roles in intracellular trafficking, including the control of endosomal morphology, protein composition, and compartment identity (HAUCKE AND DI PAOLO 2007; YANG *et al*. 2018; REDPATH *et al*. 2020; SAKURAGI AND NAGATA 2023).

Distinct lipid species residing within the bilayers of endosomes also affect the physical properties of membranes including their ability to form tubules and extrude vesicles (ANDERSEN *et al*. 2016; HIRAMA *et al*. 2017). Consistently, an increase in the proportion of certain lipid subtypes can lead to the formation of membrane microdomains within endosomes, contributing to processes that include cargo sorting (JACOBSON *et al*. 2019). Finally, specific lipids are critical for the recruitment of many key endocytic proteins that are required to carry out compartment-specific functions (GRAHAM AND KOZLOV 2010; LUCAS AND CHO 2011; KULAKOWSKI *et al*. 2018).

Glycerophospholipids are one class of membrane lipids that affect intracellular trafficking and include phosphatidylinositol (PI), phosphatidylcholine (PC), phosphatidylethanolamine (PE), and phosphatidylserine (PS) (YANG *et al*. 2018). PS is a negatively charged phospholipid and represents ∼3–10% of the overall lipid makeup of cells (YANG *et al*. 2018; KAY AND FAIRN 2019). The isolation of membrane structures *ex vivo* shows PS to be particularly abundant at the plasma membrane and in endocytic compartments, especially within recycling endosomes (LEVENTIS AND GRINSTEIN 2010). In most cells, PS is largely sequestered within the cytoplasmic leaflet of membrane bilayers (YEUNG *et al*. 2008). This asymmetry is produced and maintained by the activity of flippase proteins, which use the energy of ATP hydrolysis to transfer PS, or other types of lipids, from the non-cytosolic to the cytosolic leaflet (LEVENTIS AND GRINSTEIN 2010; ANDERSEN *et al*. 2016). The effects of PS asymmetry are two-fold (BEST *et al*. 2019). First, the shape and negative charge of PS alter membrane morphology by creating an imbalance in surface area between the two leaflets of the lipid bilayer, with the PS-enriched bilayer expanding due to a concentration of repulsive negative charges (FANANI AND AMBROGGIO 2023). Second, the negative charge of PS and its resulting alteration in membrane morphology lead to the recruitment of endocytic proteins, which may bind to PS through specific domains or may associate with PS-enriched membranes through bridging proteins (GRAHAM AND KOZLOV 2010; KULAKOWSKI *et al*. 2018). PS-associated proteins may in turn further modify compartment identity, morphology, and endocytic processes (LUCAS AND CHO 2011; UCHIDA *et al*. 2011; KAY AND GRINSTEIN 2013).

Our previous work to uncover the roles of the *C. elegans* NIMA-related kinases NEKL-2 (NEK8/9 in mammals) and NEKL-3 (NEK6/7 in mammals) in larval molting has led to the finding that NEKLs carry out conserved functions in endocytic trafficking in nematodes and mammals. In *C. elegans*, the absence of wild-type *nekl-2* or *nekl-3* leads to arrest during early larval development due to an inability to shed the old cuticle (YOCHEM *et al*. 2015; LAZETIC AND FAY 2017). Both NEKL-2 and NEKL-3 are expressed and required within hyp7, a large multinucleate epidermal syncytium that covers most of the worm midbody. Likewise, the conserved ankyrin repeat binding partners of the NEKLs, MLT-2, MLT-3, and MLT-4 (ANKS6, ANKS3, and INVS in mammals, respectively), are required for molting and function specifically within hyp7 (LAZETIC AND FAY 2017). Loss of NEKL-2 or NEKL-3 also leads to widespread defects in membrane trafficking including clathrin-dependent and -independent endocytosis, cargo recycling, and endosomal morphology (YOCHEM *et al*. 2015; LAZETIC AND FAY 2017; JOSEPH *et al*. 2020; JOSEPH *et al*. 2023).

Notably, endocytosis is required for the uptake of cargoes that are essential for the molting process, including steroid hormone precursors that are necessary for the coordinated expression of molting-related genes (YOCHEM *et al*. 1999; ASAHINA *et al*. 2000; GISSENDANNER AND SLUDER 2000; FRAND *et al*. 2005; KOUNS *et al*. 2011; KANG *et al*. 2013; LAŽETIĆ AND FAY 2017; TSIAIRIS AND GROSSHANS 2021; JOHNSON *et al*. 2023). In addition, endocytosis serves to internalize components of the old cuticle and to stabilize the surface area of the plasma membrane during periods of extensive exocytosis when new cuticular components are being secreted (JOHNSTONE AND BARRY 1996; ROBERTS *et al*. 2003; PAGE *et al*. 2006; LAŽETIĆ AND FAY 2017; MIAO *et al*. 2020).

We previously described a genetic approach to identify suppressors of *nekl*-associated molting and trafficking defects that uncovered several conserved protein phosphatases, membrane proteases, and regulators of clathrin-mediated endocytosis (JOSEPH *et al*. 2018; LAŽETIĆ *et al*. 2018; JOSEPH *et al*. 2020; JOSEPH *et al*. 2022; BINTI *et al*. 2024b). Here we report the identification of suppressing mutations in *tat-1*, which encodes the *C. elegans* ortholog of the mammalian PS flippases ATP8A1 and ATP8A2 (VAN DER MARK *et al*. 2013; ANDERSEN *et al*. 2016). We found that TAT-1 was required for the normal distribution of PS within hyp7, and that loss of TAT-1 activity led to defects in a broad range of endocytic compartments, most notably recycling endosomes marked by the *C. elegans* EHD1 ortholog, RME-1 (GRANT *et al*. 2001).

First identified in *C. elegans* as a basolateral recycling protein, Eps15 homology domain family proteins (EHD1/RME-1) are endosomal ATPases, which are thought to promote cargo recycling by remodeling membranes (GRANT *et al*. 2001; GRANT AND CAPLAN 2008). We provide evidence that TAT-1 is required for the recruitment of RME-1 to recycling endosomes via PS and by a previously identified EHD1-interacting protein, RFIP-2 (human RAB-11FIP5) (SOLINGER *et al*. 2020). In addition, depletion of either NEKl-2 or NEKL-3 led to a reduction in the abundance of RME-1– marked endosomes. Somewhat counterintuitively, however, we found that the loss of TAT-1 function may suppress *nekl*-associated defects, in part by restoring RME-1 localization to recycling endosomes in the absence of NEKL function.

## MATERIALS AND METHODS

### Strains and propagation

All *C. elegans* strains were maintained according to previously described methods and propagated at 22°C unless otherwise indicated (STIERNAGLE 2006). For the vacuolation assay, gravid hermaphrodites were placed at 25°C for ∼8 hours before being removed from plates. Progeny of these animals were then incubated at 25°C until adulthood before being imaged. Strains used in this study are listed in Table S1 in File S2.

### RNAi

Standard dsRNA injection methods were used for RNAi experiments (MONTGOMERY 2004). Primers corresponding to *tat-1*, *chat-1*, *rfip-2,* and, *rme-1 pssy-1* containing the T7 RNA polymerase binding motif were used to synthesize dsRNA as described (AHRINGER 2005). Primer information is in Table S2 in File S2. For suppression scoring,

0.5 μg/µl dsRNA was injected into adult hermaphrodites, and F1 progeny were scored for viability. For confocal imaging, 0.5 μg/µl of dsRNA was injected into adult hermaphrodites. Day-1 adult hermaphrodites from the F1 generation were placed on nematode growth medium 3× peptone agar plates containing auxin as described below, before being imaged by confocal microscopy 24 hours later.

### Reporter strain construction

Plasmids for *C. elegans* hyp7-specific expression used promoters for *semo-1/Y37A1B.5* (P_hyp7_) or *rab-5* (P*_rab-5_*) as indicated in File S1 and figure legends. Strains expressing entry vectors containing *rab-5, rab-7, tgn-38, daf-4,* and *rme-1* were created using the Gateway system (Invitrogen), pDONR221, and modified versions of the hygromycin-resistant and miniMos-enabled vector pCFJ1662 (Addgene #51482; gifts from Erik Jorgensen and Barth Grant; (JOSEPH *et al*. 2023). The *nekl-3* promoter (P_nekl-3_) was transferred into pCFJ1662 vector by restriction digest (pDF477). An entry vector containing mouse *mfge8*/*Lact^C1C2^* coding region (Addgene #52987) (MAPES *et al*. 2012) was transferred into the pDF477 destination vector using the pENTR/dTOPO cloning system (Thermo Fisher Scientific). Single-copy integrations were obtained using miniMos procedures (FROKJAER-JENSEN *et al*. 2014).

### CRISPR mutant alleles

Loss-of-function alleles of *tat-1* (E853K CRISPR phenocopy, E170Q, S1069stop) were created using established CRISPR-Cas9 protocols (FARBOUD AND MEYER 2015; FARBOUD *et al*. 2019; GHANTA *et al*. 2021). sgRNA and repair templates were synthesized by Integrated DNA Technologies and Dharmacon-Horizon Discovery; Ape, and CRISPRcruncher were used in the design of guideRNA and repair templates (FAY *et al*. 2021; DAVIS AND JORGENSEN 2022). Primer, sgRNA, and repair template sequences are provided in Table S3 in File S2.

### Image acquisition and analysis

Confocal fluorescence images were acquired using cellSense (ver 4.2) (Olympus Corporation) on an Olympus IX83 inverted microscope and Yokogawa spinning disc confocal head (CSU-W1). z-Stack images were acquired using a 100×, 1.35 N.A. silicone oil objective. Quantification of mean intensity (measured in arbitrary units, a.u.), average area of vesicles (measured as area in micrometers, µm^2^), colocalization analysis, and vacuolation assay were performed using Fiji (SCHINDELIN *et al*. 2012) and Olympus cellSens software. To quantify mean intensity, for a z-plane of interest, the intensity of the background of an image was measured using the rectangle selection tool in a region of the image in which no fluorescence was visible. This value was subtracted from the intensity value obtained from the hyp7 of each animal in each picture by using the polygon selection tool to select the appropriate region (JOSEPH *et al*. 2023).

To quantify the average area of vesicles for a z-plane of interest, either rolling ball background subtraction (radius = 50 pixels, figs) or the minimum filter (radius = 10 pixels, figs) was applied to the raw images using Fiji. If the minimum filter was used, the filtered image was subtracted from the raw image using the image calculator function.

Processed images were then thresholded, and the Despeckle function was applied to remove signal noise of ≤1 pixel in size. The Analyze Particles function was applied to determine the average area of vesicles. Within each experiment, the same threshold algorithm was used for all images (JOSEPH *et al*. 2023).

For quantifying colocalization, the raw z-stack images were deconvoluted using the Wiener deconvolution algorithm available in cellSens (ver. 4.2); spectral unmixing was also applied to GFP::RAB-11; CHAT-1::mKate images (Fig. 4a–c’). The appropriate z-plane was extracted from the raw and deconvoluted images. The deconvoluted image was subsequently thresholded to obtain a binary image to be used as a mask. The binary mask was then combined with the raw image using the “AND” Boolean operation. hyp7 was selected in the combined images using the polygon tool, and the BIOP JACoP plugin was used to calculate the Pearson’s correlation coefficient (R) and Manders’ overlap (M) (BOLTE AND CORDELIERES 2006). Combined images were used to determine cases of significant overlap in contrast to random co-occurrence.

From the combined images, a 100x100 µm square (10,000 µm^2^) was sampled, and M and R values were calculated using the BIOP JACop plugin (“inset”). The red channel was rotated ninety degrees in relation to the green channel using the transform function, thus creating a random distribution of green and red pixels of interest; M and R values were subsequently calculated using the BIOP JACoP plugin (“rotated”) (DUNN *et al*. 2011).

To quantify the presence of vacuoles in the intestine or epidermis, a z-stack of images was acquired in young adults. Any vacuole that measured 5 µm or larger in diameter using the line tool (Fiji) was counted.

### Auxin treatment

Depletion of auxin-inducible degron (AID)-tagged NEKL alleles was carried out as described (JOSEPH *et al*. 2020; JOSEPH *et al*. 2023). Briefly, a stock solution of auxin (indole-3-acetic acid) was created by dissolving 0.7 g of auxin (purchased from Alfa Aesar) in 10 ml of 100% ethanol. A mixture of 25 μl auxin stock solution and 225 μl autoclaved deionized water was then spread onto each standard agar plate. Larval stage 4 (L4) worms were grown for 24 hours on standard NGM plates and were then transferred to the agar plates containing auxin (indole-3-acetic acid).

### Protein structure analysis

Predicted protein structures were generated using AlphaFold2 or AlphaFold3 and UCSF ChimeraX 1.7 and structure prediction software (JUMPER *et al*. 2021; PETTERSEN *et al*. 2021; EVANS 2022; ABRAMSON *et al*. 2024).

### Statistical analysis

Statistical tests were performed with GraphPad Prism software using standard methods (FAY AND GEROW 2013).

## RESULTS

### Mutations in *tat-1* suppress *nekl* loss-of-function molting defects

We previously described a forward genetic screen and whole-genome sequencing pipeline to identify mutations that suppress the larval lethality of *nekl* loss-of-function mutations (JOSEPH *et al*. 2018). The screen makes use of weak hypomorphic alleles of *nekl-2* (*fd81*) and *nekl-3* (*gk894345*), which are aphenotypic as single mutants but when combined are synthetically lethal, with only 2.8% of *nekl-2(fd81); nekl-3*(*gk894345*) double mutants (hereafter *nekl-2; nekl-3*) reaching adulthood (Fig. 1a, c) (LAŽETIĆ AND FAY 2017). Double mutants are maintained by a GFP-marked rescuing extrachromosomal array containing wild-type copies of *nekl-3* (Fig. 1a). After mutagenesis, strains that have acquired an extragenic suppressor are identified by their viability in the absence of the rescuing array (JOSEPH *et al*. 2018).

**Fig. 1.**
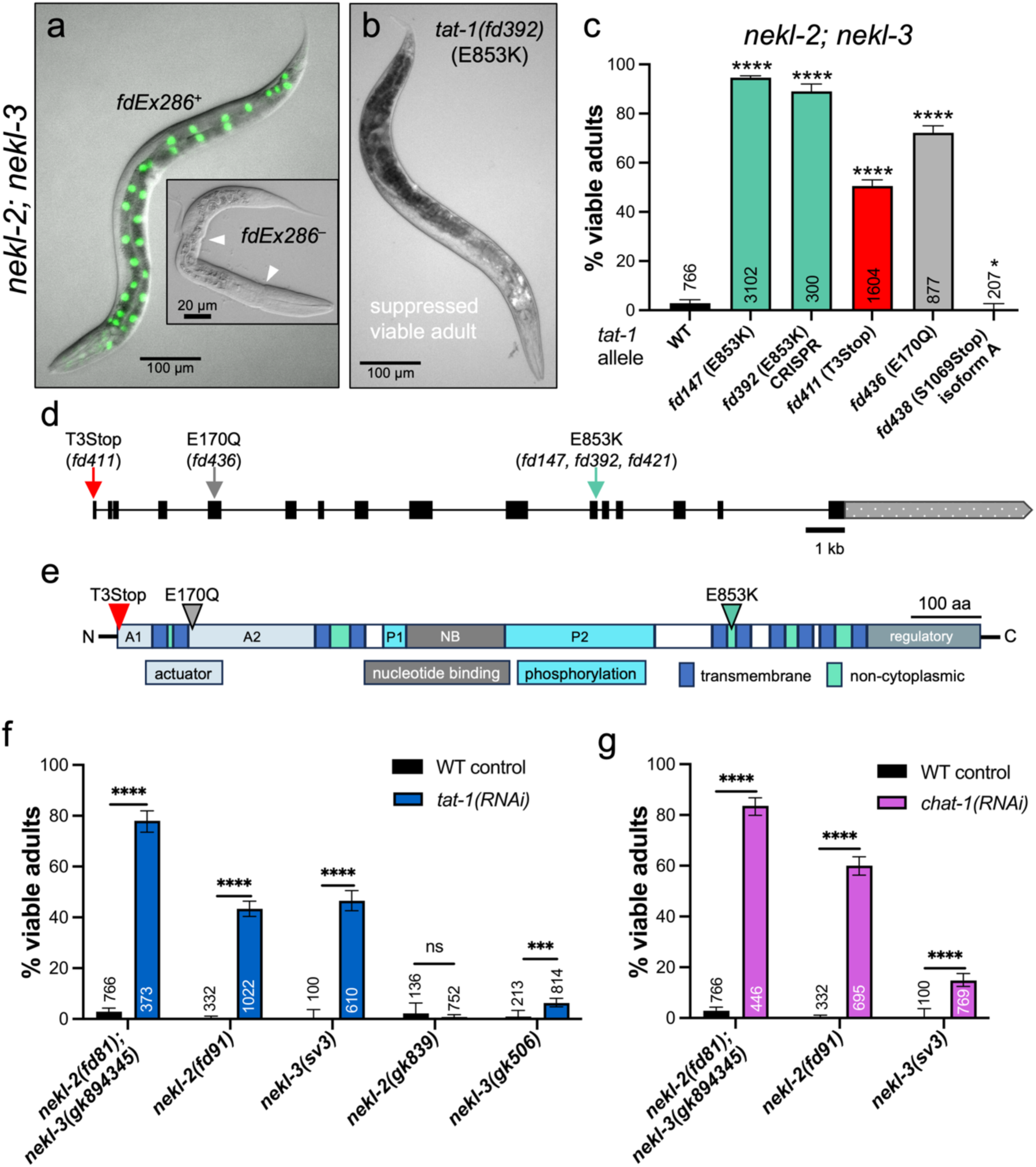
Suppression of *nekl* molting defects by *tat-1* and *chat-1*. (a) *nekl-2(fd81); nekl-3(gk894345)* double mutants arrest as larvae (inset) but are rescued by an extrachromosomal array (*fdEx286*) containing wild-type copies of *nekl-3* and a broadly expressed GFP reporter (*sur-5::GFP*). White arrowheads (inset) indicate anterior and posterior boundaries where the old cuticle has remained attached to the midbody. (b) Example of a suppressed *nekl-2(fd81); tat-1(fd147); nekl-3(gk894345)* adult. (c) Bar graph showing the percentage of viable adult (non-molting-defective) progeny for the indicated genotypes and their corresponding protein-based changes. (d,e) Schematic of (d) *tat-1* genomic locus and (e) TAT-1 protein; arrows/arrowheads indicate the locations of alleles. A1 and A2 indicate the bipartite actuator domain; P1 and P2, the bipartite phosphorylation domain; NB, the nucleotide (ATP) binding domain. (f, g) RNAi knockdown of (f) *tat-1* and (g) *chat-1* by microinjection of dsRNA robustly suppresses *nekl-2(fd81); nekl-3(gk894345)* double mutants and moderate loss-of-function alleles of *nekl-2* (*fd91*) and *nekl-3* (*sv3*) but not presumed null alleles (*gk839* and *gk506*). Note that the data for control *nekl-2; nekl-3* in panels c, f, and g are identical. All graphs show the mean and 95% confidence interval (CI; error bars). Statistical significance was determined by using Fisher’s exact test; ****p ≤ 0.001, ***p ≤ 0.001, *p ≤ 0.05; ns, not significant. Raw data for all panels are available in Supplementary File 1.

From our earlier suppressor screen, we isolated *fd147*, a recessive allele that enables ∼95% of *nekl-2; nekl-3* animals to reach adulthood (JOSEPH *et al*. 2018). Whole-genome sequencing identified mutations in three coding regions including a G-to-A transversion in the eleventh exon of *tat-1* (Fig. 1d), leading to the substitution of lysine for a glutamic acid at amino acid 853 (Fig. 1d, e). CRISPR phenocopy of *tat-1*(E853K; *fd392*) led to ∼90% of *nekl-2; nekl-3* double mutants reaching adulthood, indicating that the *fd147* causal mutation resides in *tat-1* (Fig. 1c). Consistent with *tat-1*(E853K) being a reduction-of-function allele, *tat-1(RNAi)* (using dsRNA injection methods) led to strong suppression of *nekl-2; nekl-3* mutants (Fig. 1f). We also used CRIPSR to generate a putative null allele of *tat-1* containing several stop codons and a frameshift following the second amino acid of TAT-1 (T3STOP (*fd411)*; Fig. 1c). Whereas clear suppression of *nekl-2; nekl-3* mutants was conferred by this allele, it was notably weaker (∼50%) than that with either *tat-1*(E853K) or *tat-1(RNAi)* (Fig. 1c, f), suggesting that hypomorphic alleles of *tat-1* may be better able to suppress *nekl-2; nekl-3* molting defects than a complete loss of TAT-1 function.

TAT-1 is a member of the P4 family of P-type ATPases, which use ATP hydrolysis to translocate lipids between the inner and outer leaflets of membrane bilayers (DARLAND-RANSOM *et al*. 2008; RUAUD *et al*. 2009). TAT-1 is most similar to the human class 1a P4-ATPases ATP8A1 and ATP8A2, which function primarily as PS flippases (ANDERSEN *et al*. 2016). TAT-1 is predicted to contain 10 transmembrane (TM) domains, resulting in five non-cytosolic loops and six cytoplasmic domains, in addition to internal domains at the N and C termini (JUMPER *et al*. 2021). Based on homology, TM1–6 are predicted to be the main path by which phospholipids are transported across membranes, whereas the adjacent non-cytosolic loops have been implicated in the binding of lipids prior to translocation (DUAN AND LI 2024). The E853K mutation falls within the third extracellular loop between the fifth and sixth TM domain and may therefore affect PS recognition and translocation. Notably, E853 is 100% conserved in mammals and resides in a region of TAT-1 marked by acid surface residues (Fig S1a– a’’). TAT-1 contains four discontinuous cytoplasmic domains found in all P4-ATPases including a central nucleotide-binding domain, a bipartite phosphorylation domain (P1 and P2) that is predicted to undergo modification at a conserved aspartic acid residue (D454)(CHEN *et al*. 2019), a bipartite N-terminal actuator domain (A1 and A2) required for the hydrolysis of aspartic acid and completion of the catalytic cycle, and a proposed C-terminal regulatory domain that may direct subcellular localization and enzymatic activity (Fig. 1e; Fig. S1a, a’) (ANDERSEN *et al*. 2016). To determine if a reduction of TAT-1 catalytic activity is sufficient for *nekl* suppression, we used CRISPR to change a glutamic acid residue within the actuator domain (DG**E**T motif) to glutamine (*fd436* E170Q, Fig. 1c–e), a substitution that inhibits hydrolysis of the aspartyl phosphoryl bond in mammalian TAT-1 orthologs (COLEMAN *et al*. 2012). Notably, E170Q suppressed *nekl* molting defects at levels that were intermediate between the E853K and T3STOP mutants, consistent with *nekl* suppression resulting from a reduction in TAT-1 flippase activity (Fig. 1c–e).

Data available on WormBase indicate that *tat-1* has three isoforms, which differ specifically at their C termini, with *tat-1a* and *tat-1c* being the most abundant isoforms (Fig. S1b) (STERNBERG *et al*. 2024). To test the relative importance of *tat-1a* and *tat-1c* isoforms in *nekl* suppression, we used CRISPR to generate an early stop codon at the beginning of the sixteenth exon of *tat-1a* (S1069Stop; *fd438*), thus preventing translation of the last two exons of *tat-1a* while leaving the *tat-1b* and *tat-1c* isoforms intact. The S1069Stop mutation failed to suppress *nekl-2; nekl-3* mutants (Fig. 1c), suggesting that loss of TAT-1C may be most critical for *nekl* suppression or that *tat-1* isoforms may be functionally redundant.

To test whether suppression of *nekl-2; nekl-3* defects by loss of TAT-1 is specific to the function of either *nekl-2* or *nekl-3*, we injected *tat-1* dsRNA into strains containing individual moderate or strong loss-of-function alleles of *nekl-2* or *nekl-3*. Whereas we observed substantial suppression of *nekl-2(fd91)* and *nekl-3(sv3)*, which are moderate loss-of-function alleles, we detected only weak suppression of *nekl-3(gk506)* null mutants and no suppression of *nekl-2(gk839)* null mutants (Fig. 1f). These results demonstrate that *tat-1*–based suppression is not specific to either *nekl-2* or *nekl-3* functions and place *tat-1* among the stronger *nekl* suppressors identified to date (JOSEPH *et al*. 2018).

### TAT-1 colocalizes with CHAT-1 and is dependent on CHAT-1 for its localization

Most mammalian P4-ATPases form a heteromeric complex with a CDC50/LEM3 family β-subunit, which is essential for their proper folding, catalytic activity, and exit from the endoplasmic reticulum (KATOH AND KATOH 2004; COLEMAN AND MOLDAY 2011; ANDERSEN *et al*. 2016). In *C. elegans*, CHAT-1 (<u>ch</u>aperone of *tat-1*) has been identified as the ortholog of CDC50A, the mammalian β-subunit of ATP8A1 and ATP8A2 (CHEN *et al*. 2010). In contrast to the documented α–β contact site between ATP8A2 and CDC50A, AlphaFold modeling places the two TM domains of CHAT-1 adjacent to TM7 and TM10 of TAT-1 (Fig. S1c–c’’), more closely matching the reported configuration of P-type ATPase family Na^+^/K^+^-ATPase α- and β-subunits (ANDERSEN *et al*. 2016). In this orientation, the non-cytosolic portion of CHAT-1 would be predicted to interact directly with the non-cytosolic loops of TAT-1 (Fig. S1c–c’’), suggesting a possible direct role for CHAT-1 in phospholipid binding and flippase activity, consistent with previous reports (LENOIR *et al*. 2009; COLEMAN AND MOLDAY 2011). Interestingly, E853 of TAT-1 is proximal to a region of TAT-1 that interacts with CHAT-1 (Fig. S1a’’, c–c’’), suggesting that E853K could impact the interaction of TAT-1 with CHAT-1. As anticipated, when *nekl-2; nekl-3* animals were injected with *chat-1* dsRNA, suppression was detected at levels equivalent to those observed for *tat-1(RNAi)* (Fig. 1f, g). Furthermore, like *tat-1(RNAi)*, suppression of moderate loss-of-function alleles of *nekl-2* and *nekl-3* was observed after *chat-1(RNAi)* (Fig. 1g).

Our genetic and protein modeling data, as well as prior studies on TAT-1 and CHAT-1 family members, predict that TAT-1 and CHAT-1 colocalize in vivo (BRYDE *et al*. 2010; CHEN *et al*. 2010). To test this, we obtained fluorescently tagged alleles of TAT-1 (GFP and mScarlet) and CHAT-1 (mKate) using CRISPR methods (SunyBiotech) and examined their subcellular localization patterns in the epidermis (Fig. 2a, b), where NEKL-2 and NEKL-3 are expressed and specifically required for molting (YOCHEM *et al*. 2015; LAZETIC AND FAY 2017). TAT-1::GFP was distributed in a mesh-like or punctate localization pattern throughout the apical portion of hyp7, a large multi-nucleate epidermal syncytium covering most of the worm midbody (SCHROEDER AND HALL 2021), but it was excluded from the adjacent epidermal seam cell (Fig. 2c, c’; Fig. S1d, d’). Likewise, worms expressing CHAT-1::mKate displayed a similar localization pattern within hyp7 (Fig 2d, d’; Fig. S1e, e’), and animals expressing both fluorescently tagged proteins showed extensive overlap, as supported by Manders’ overlap and Pearson’s correlation coefficients (Fig. 2e–g). We note that for these and other co-localization experiments we measured correlation coefficients in the entire section of hyp7 that was visible within each frame, as well as within representative square insets. The 90° rotation of each inset served as a co-localization control for our studies (File S1) (DUNN *et al*. 2011).

**Fig. 2.**
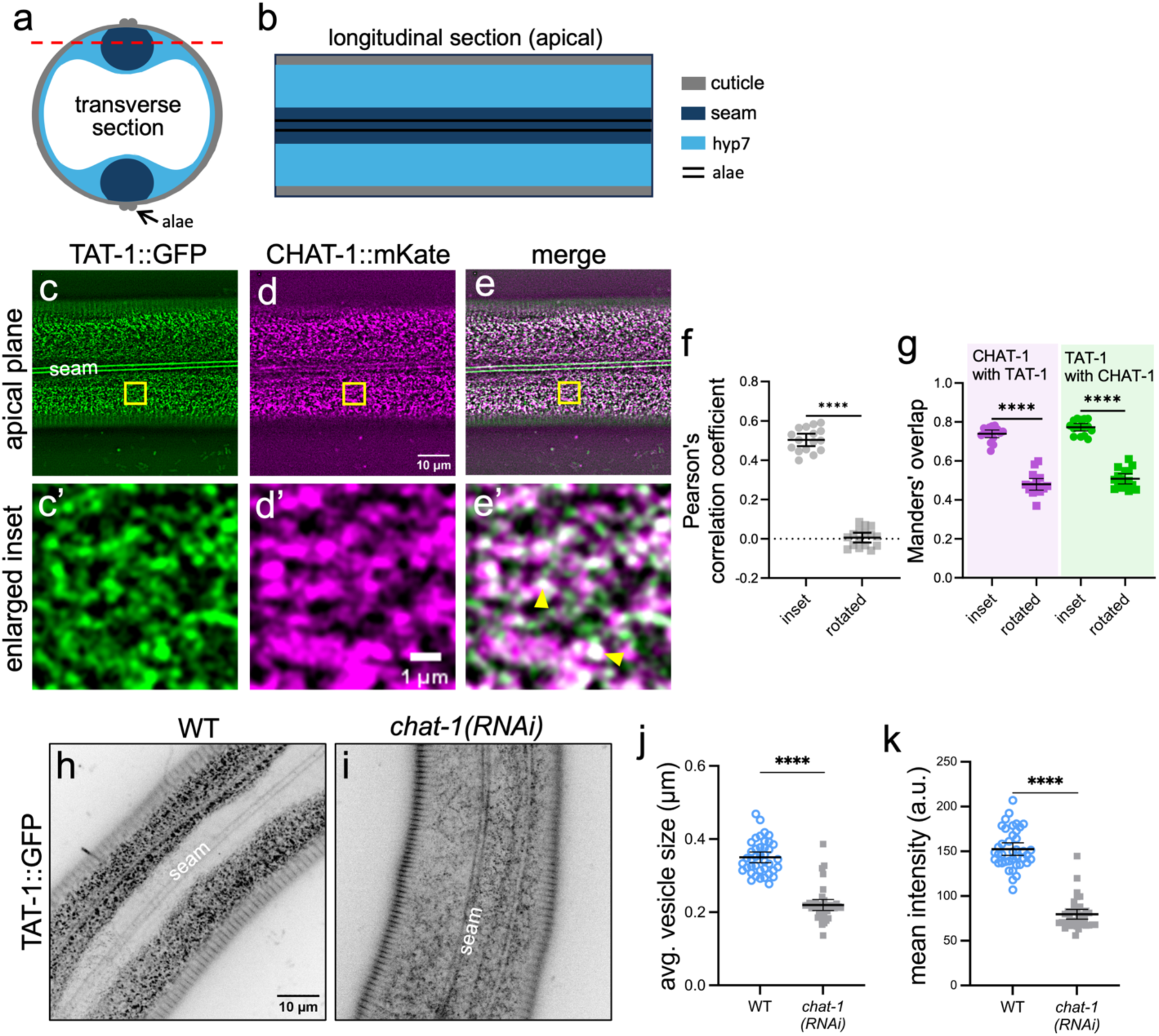
CHAT-1 regulates TAT-1 epidermal abundance. (a, b) Schematic representation of the adult *C. elegans* epidermis viewed (a) as a transverse cross-section and (b) from a longitudinal apical plane just underneath the cuticle. Red dashed line in (a) indicates the approximate location of the apical plane in (b). Expression and localization analyses were carried out for hyp7 specifically and excluded the seam cell. (c–e’) Representative confocal images of day-1 adult hermaphrodites expressing (c, c’) TAT-1::GFP, (d, d’) and CHAT-1A::mKate, along with both markers (e, e’) (merge). Yellow arrowheads in (e’) indicate examples of colocalization in white. Yellow boxes in (c–e) indicate the enlarged insets in (c’–e’). (f, g) Dot plots of (f) Pearson’s correlation coefficient and (g) Manders’ overlap between TAT-1::GFP and CHAT-1A::mKate (n = 35 animals). Rotated indicates 90° rotation of one of the two inset regions and serves as a control; also see Materials and Methods. (h–k) TAT-1::GFP expression in representative control (wild type, WT) and (i) *chat-1(RNAi)* animals, along with corresponding dot plots that show (j) the average area of vesicles and (k) the mean intensities in these animals (n ≥ 39 animals for each experiment). The reduced visibility of the seam cell in is due to the lower focal plane captured in this image versus in (h), reflecting a wider apical–basal dispersion of TAT-1::GFP in *chat-1(RNAi)* animals. All plots show the mean and 95% CI. Statistical significance was determined using an unpaired t-test; ****p ≤ 0.0001. Raw data for all panels are available in Supplementary File 1.

Based on prior work, along with our observation that *chat-1*(*RNAi*) suppresses *nekl-2; nekl-3* molting defects, we predicted that knockdown of *chat-1* would alter TAT-1::GFP localization in the epidermis (COLEMAN AND MOLDAY 2011). Notably, *chat-1(RNAi)* led to a decrease in the average area of TAT-1::GFP vesicles and to a decrease in the mean intensity of TAT-1::GFP (Fig. 2h–k). Moreover, *chat-1(RNAi)* led to poorly defined TAT-1::GFP puncta and a diffuse fluorescent signal that appeared to be evenly distributed throughout the apical and basal layers of hyp7. Our results indicate that CHAT-1 is required for TAT-1 localization and abundance within the epidermis, although it remains possible that CHAT-1 may play additional roles in PS binding and flipping.

### Loss of TAT-1 affects multiple endosomal compartments

Previous studies on *C. elegans* and mammalian TAT-1 family members have implicated TAT-1 in multiple steps of membrane trafficking including endocytosis, exocytosis, autophagy, cell engulfment, and transport through the endocytic system (DARLAND-RANSOM *et al*. 2008; RUAUD *et al*. 2009; CHEN *et al*. 2010; NILSSON *et al*. 2011; MAPES *et al*. 2012; RAIDERS *et al*. 2021). This includes reports indicating that cytoplasmic-facing PS on the plasma membrane plays a role in promoting the budding and pinching-off of clathrin-coated vesicles by recruiting F-BAR domain– containing proteins, which support membrane bending and scission (ITOH AND DE CAMILLI 2006; LEMMON 2008; VARGA *et al*. 2020). To test whether inhibition of TAT-1 affects clathrin-mediated endocytosis in the *C. elegans* epidermis, we introduced *tat-1*(E853K) into strains containing fluorescently tagged clathrin light (CLIC-1) and heavy (CHC-1) chains and looked for alterations in hyp7 clathrin localization. In wild type, mScarlet::CLIC-1 and GFP::CHC-1 formed small, regularly dispersed apical puncta throughout hyp7. These clathrin markers showed a modest increase in mean intensity in *tat-1*(E853K) mutants, although the average area of vesicles remained indistinguishable from wild type (Fig. S2a–h), suggesting that TAT-1 may play a relatively minor role in the internalization of clathrin-coated vesicles in the epidermis.

To assess the effects of TAT-1 inhibition on endosomal compartments we used strains expressing fluorescently tagged RAB proteins, a large and highly conserved family of small GTPases that function as endosome-specific organizers (ZERIAL AND MCBRIDE 2001; GROSSHANS *et al*. 2006; CHUA AND TANG 2018; HOMMA *et al*. 2021). In contrast to clathrin, we observed more-pronounced effects on endosomal compartments in *tat-1*(E853K) mutants including changes in early (GFP::RAB-5), late (GFP::RAB-7), and recycling (RAB-10, RAB-11, RME-1) endosomes (Fig. 3; Fig. S2i–l). In the case of RAB-marked endosomal compartments, we observed a uniform increase in the average mean intensity of markers ranging from ∼1.5-fold (GFP::RAB-11 and GFP::RAB-7) to ∼1.2-fold (GFP::RAB-10 and GFP::RAB-5). In addition, *tat-1*(E853K) led to an increase in the average area of RAB-expressing vesicles/puncta, most notably for GFP::RAB-7 (∼1.5-fold). Enlarged GFP::RAB-5 and GFP::RAB-7 compartments in *tat-1* mutants were also positioned in more basal regions of hyp7 (Fig. 3c, e). In addition, we observed fewer GFP::RAB-7–marked tubules in *tat-1* mutants, as compared to wild type (Fig. 3d, e). These results indicate that TAT-1 may restrict the localization of the RAB proteins or may control the size and morphologies of their associated endosomal compartments. These findings are consistent with previous studies showing that RAB protein localization in the intestine is also affected by loss of TAT-1 (RUAUD *et al*. 2009; CHEN *et al*. 2010; LI *et al*. 2013).

**Fig. 3.**
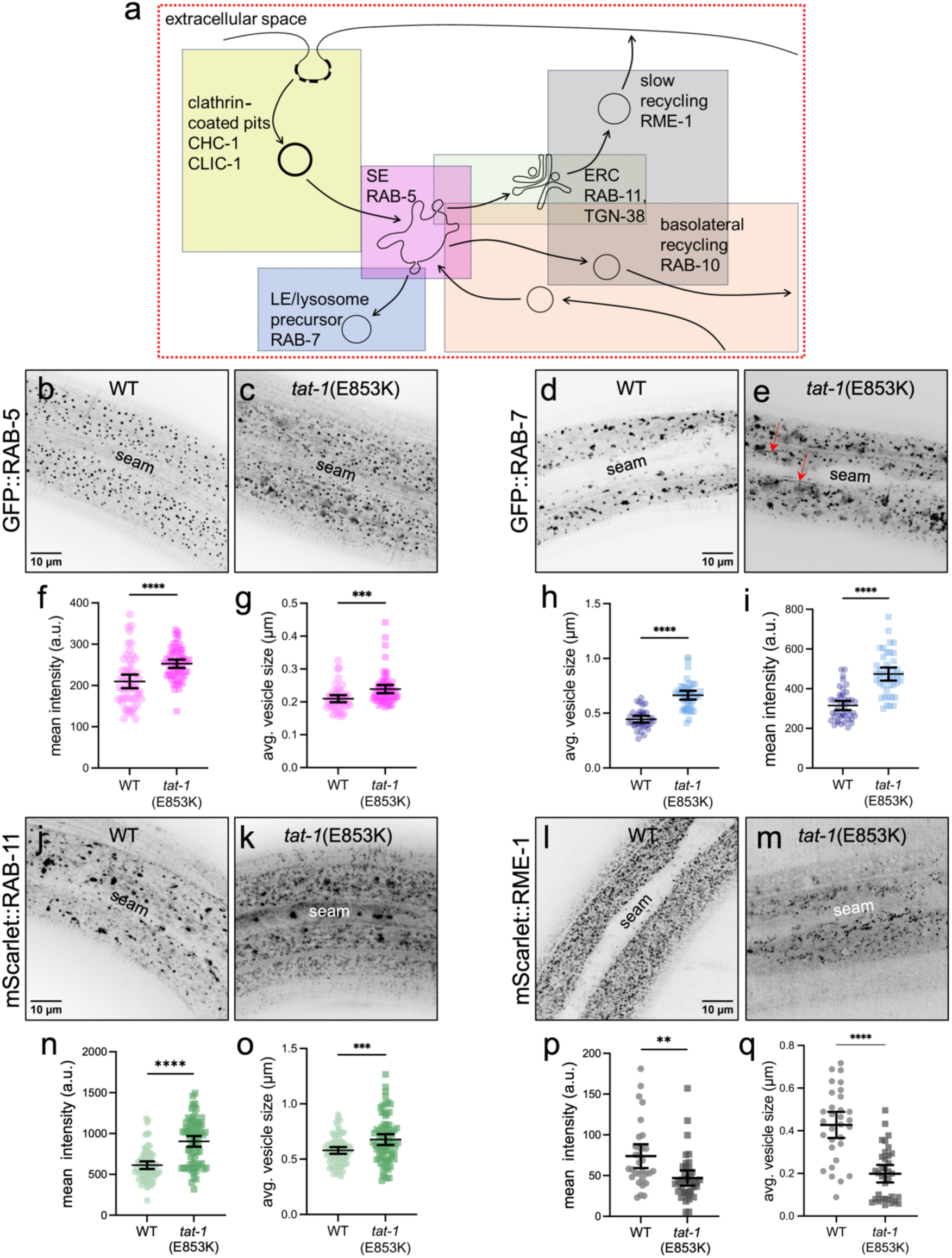
Loss of TAT-1 function alters endosome morphology. (a) Schematic of endocytosis and endosomal compartments. Shaded boxes outline approximate location(s) of endocytic proteins and compartments within the epidermis. SE, sorting (early) endosome; LE, late endosome; ERC, endocytic recycling compartment. (b–e and j–m) Representative confocal images of WT and *tat-1*(E853K) day-1 adults expressing either (b, c) P*_rab-5_*::GFP::RAB-5, (d, e) P_hyp7_::GFP::RAB-7, (j, k) mScarlet::RAB-11, or (l, m) P_hyp7_::mScarlet::RME-1. Red arrows in (e) indicate elongated tubule-like GFP::RAB-7–marked compartments. (f–i and n–q) Dot plots showing the (f, h, n, p) mean intensity and (g, i, o, q) average vesicle area of corresponding genotypes. All plots show the mean and 95% CI. Statistical significance was determined using an unpaired t-test; ****p ≤ 0.0001, **p ≤ 0.01. The number of worms analyzed is indicated. Raw data for all panels are available in Supplementary File 1.

**Fig. 4.**
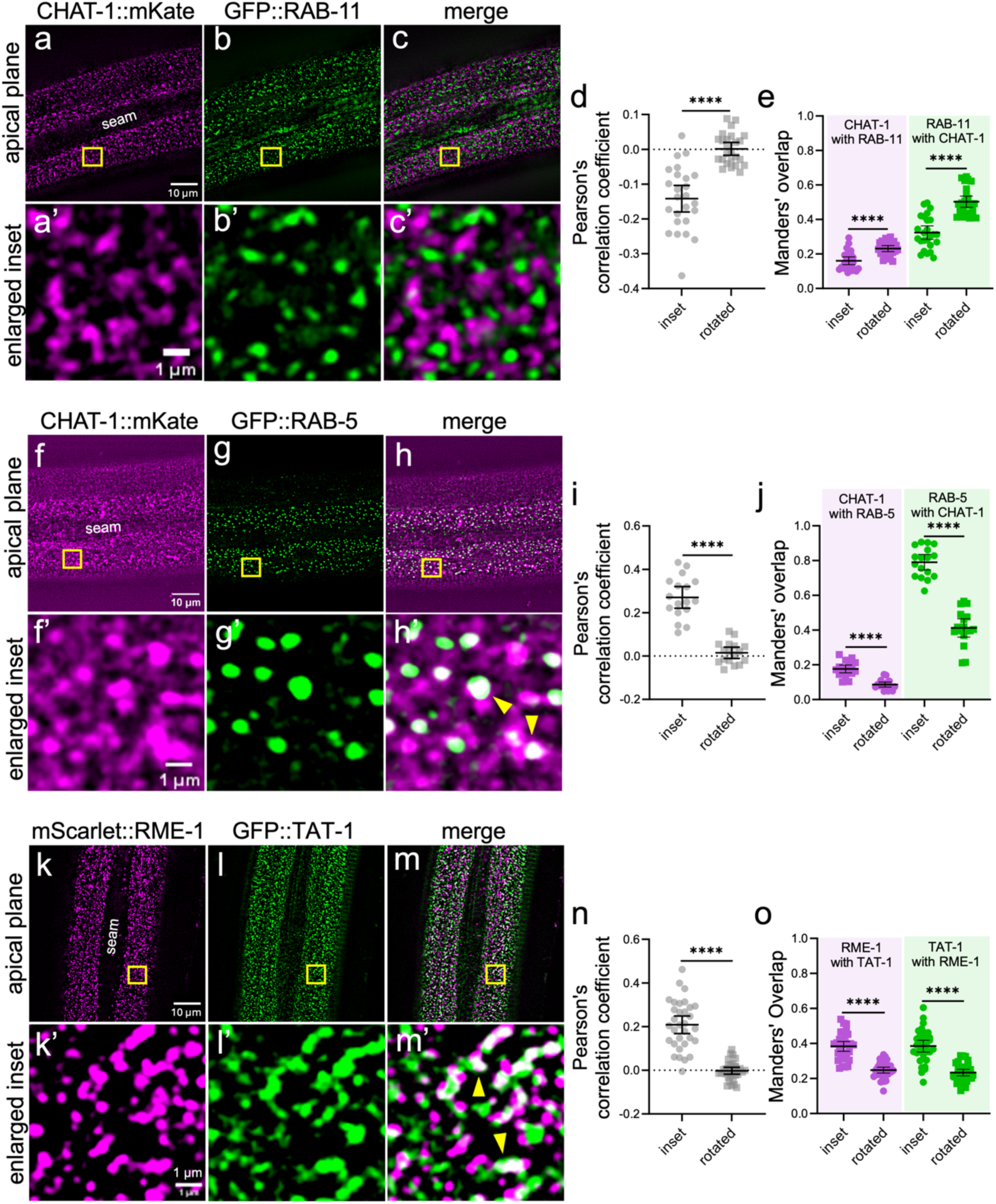
TAT-1 colocalizes to endocytic compartments. (a–c’, f–h’, k–m’) Representative confocal images of day-1 adults showing CHAT-1::mKate localization in the epidermis relative to (a–c’) GFP::RAB-11 (n = 25) and (f–h’) P_rab-5_::GFP::RAB-5 (n = 17), along with (k–m’) TAT-1::GFP relative to P_hyp7_::mScarlet::RME-1 (n = 32). Yellow boxes indicate enlarged insets; yellow arrowheads indicate examples of overlap in white. (d, e, i, j, n, o) Colocalization was quantified using (d, i, n) Pearson’s correlation coefficient and (e, j, o) Manders’ overlap. Dot plots show the mean and 95% CI. Statistical significance between rotated and inset values was determined by unpaired t-test; ****p ≤ 0.0001. Raw data for all panels are available in Supplementary File 1.

In contrast to the RAB markers, loss of *tat-1* activity led to a ∼1.6-fold decrease in the mean intensity of mScarlet::RME-1 and to a ∼2.2-fold decrease in the average area of vesicles/puncta. This finding suggests that, unlike the RAB proteins examined, TAT-1 plays a positive role in the regulation of RME-1, which localizes to the apical region of the *C. elegans* epidermis at what are predicted to be recycling endosomes. This result is consistent with a previous study in mammalian cells implicating ATPA81/2 in the localization of the mammalian RME-1 homolog, EHD1 (LEE *et al*. 2015). Furthermore, previous studies looking at the *C. elegans* intestine reported that loss of TAT-1 and CHAT-1 leads to a reduction in RME-1–marked compartments at basolateral membranes (RUAUD *et al*. 2009; CHEN *et al*. 2010; LI *et al*. 2013).

Collectively, our findings implicate TAT-1 in the maintenance of multiple membrane trafficking compartments including positive and negative roles within endosomes.

Previous work has described a temperature-sensitive phenotype resulting from a loss of *tat-1* function that leads to the abnormal accumulation of vacuoles within the intestines of worms grown at 25°C (RUAUD *et al*. 2009). These vacuoles are believed to be enlarged endosomes co-labeled by what are typically mutually exclusive compartment markers including hybrid vesicles expressing RAB-5–RAB-7 and RAB-10–RME-1 (CHEN *et al*. 2010; LI *et al*. 2013). Consistent with this report, we observed an increase in the proportion of *tat-1*(E853K) adults with vacuolated intestines at 25°C (∼34%) versus wild-type worms (7%) (Fig. S3). Moreover, although we observed vacuoles in the epidermis of *tat-1*(E853K) mutants grown at 25°C (26%), this percentage was not significantly higher than that seen in wild-type worms at 25°C (18%) (Fig. S3a–d), nor did we detect epidermal vacuoles in *tat-1*(E853K) or wild-type worms at 22°C.

Nevertheless, given our finding that *tat-1*(E853K) mutants showed variably enlarged compartments marked with RAB-5, RAB-7, RAB-10, and RAB-11, we were curious to test a subset of compartment markers for colocalization to determine if hybrid compartments might form in the epidermis after *tat-1(RNAi)*. In wild-type worms, we observed little or no co-localization between RAB-5–RME-1 and RAB-11–RME-1 and minimal co-localization between RAB-5–RAB-7 and RAB-5–RAB-11 (Fig. S3e–l, S4). Furthermore, we failed to observe an increase in the extent of overlap in doubly marked RAB-5–RAB-7, RAB-5–RAB-11, RAB-5–RME-1, and RAB-11–RME-1 worms treated with *tat-1(RNAi)* versus wild-type controls (Fig. S3e–l, S4). Therefore, in contrast to results reported for compartment markers and enlarged vacuoles in the intestine, we did not find evidence for the formation of hybrid compartments in the epidermis after TAT-1 inhibition.

### TAT-1 localizes to RAB-5–early and RME-1–recycling endosomes in the epidermis

Given that loss of TAT-1 led to effects in multiple membrane trafficking compartments, we wanted to determine the localization pattern of TAT-1. We thus examined co-localization of TAT-1::GFP or CHAT-1::mKate with the compartment markers that were altered after reduction of TAT-1 activity. We note that CHAT-1::mKate was used as a red marker in place of a TAT-1::mScarlet fusion, which exhibited non-specific localization to lysosomes, as has been observed for some fusion proteins marked by dsRed family fluorophores (CLANCY *et al*. 2023).

Although we observed modest effects on clathrin markers after *tat-1* inhibition, no overlap was detected between a clathrin light chain fusion protein (mScarlet::CLIC-1) and TAT-1::GFP (Fig. S5a–e), suggesting that the effects of TAT-1 loss on clathrin-mediated endocytosis are likely indirect. Likewise, we failed to detect colocalization of TAT-1::GFP with the endosomal markers RFP::RAB-7 and mScarlet::RAB-10 (Fig. S5f– o), again implying that effects on these compartments and proteins may be indirect.

Interestingly, we observed a strong negative correlation in the localization of GFP::RAB-11 and CHAT-1::mKate, leading to a negative Pearson’s correlation coefficient and an increase in Manders’ overlap in the rotated inset controls (Fig. 4a–e). This finding suggests that TAT-1 and RAB-11 localize to discrete, mutually exclusive compartments and that TAT-1 and/or RAB-11 may restrict each other’s binding.

In contrast, we observed strong colocalization between CHAT-1::mKate and GFP::RAB-5 and between TAT-1::GFP and mScarlet::RME-1 (Fig. 4f–o). Specifically, CHAT-1 was detected in a large proportion of GFP::RAB-5–marked round puncta (Manders’ overlap = ∼0.8) (Fig. 4f–j), suggesting that CHAT-1–TAT-1 are present on the membrane of early endosomes, where they may limit RAB-5 localization during the process of endosomal maturation. TAT-1::GFP also strongly overlapped with the mesh-like pattern of mScarlet::RME-1 in the apical-most portion of hyp7 (Fig. 4k–o), consistent with TAT-1 directly promoting the accumulation of RME-1 on apical recycling endosomes (Fig. 4).

### A PS biosensor that co-localizes with CHAT-1 and RME-1 is altered by TAT-1 inhibition

Our results thus far support a model in which TAT-1 promotes the accumulation of PS on the cytoplasmic leaflet of apical recycling endosomes (Fig. 6a), leading to the recruitment of RME-1. To test this model, we used mouse Lactadherin^C1C2^ (mfge8), a PS-binding protein fragment that binds PS in *C. elegans* and other systems (ANDERSEN *et al*. 2000; DARLAND-RANSOM *et al*. 2008; MAPES *et al*. 2012; DEL VECCHIO AND STAHELIN 2018). Specifically, wke integrated a single-copy mNeonGreen::Lact^C1C2^ transgene expressed under the control of the *nekl-3* promoter to drive hyp7-specific sensor expression at low-to-moderate levels (FROKJAER-JENSEN *et al*. 2014). As anticipated, P*_nekl-3_*::mNeonGreen::Lact^C1C2^ was detected in the apical region of hyp7 and closely resembled the mesh-like localization pattern observed in fluorescently tagged TAT-1, CHAT-1, and RME-1 strains (Fig. S1f, f’). Moreover, both CHAT-1::mKate and mScarlet::RME-1 displayed strong colocalization with P*_nekl-3_*::mNeonGreen ::Lact^C1C2^, based on Manders’ overlap and Pearson’s correlation coefficients (Fig. 5a–e, j–n). This finding indicates that RME-1 binds to endosomes that are enriched for cytoplasmic-facing PS and is consistent with TAT-1–CHAT-1 promoting the translocation of PS to the cytoplasmic surface of endosomal membranes.

**Fig. 5.**
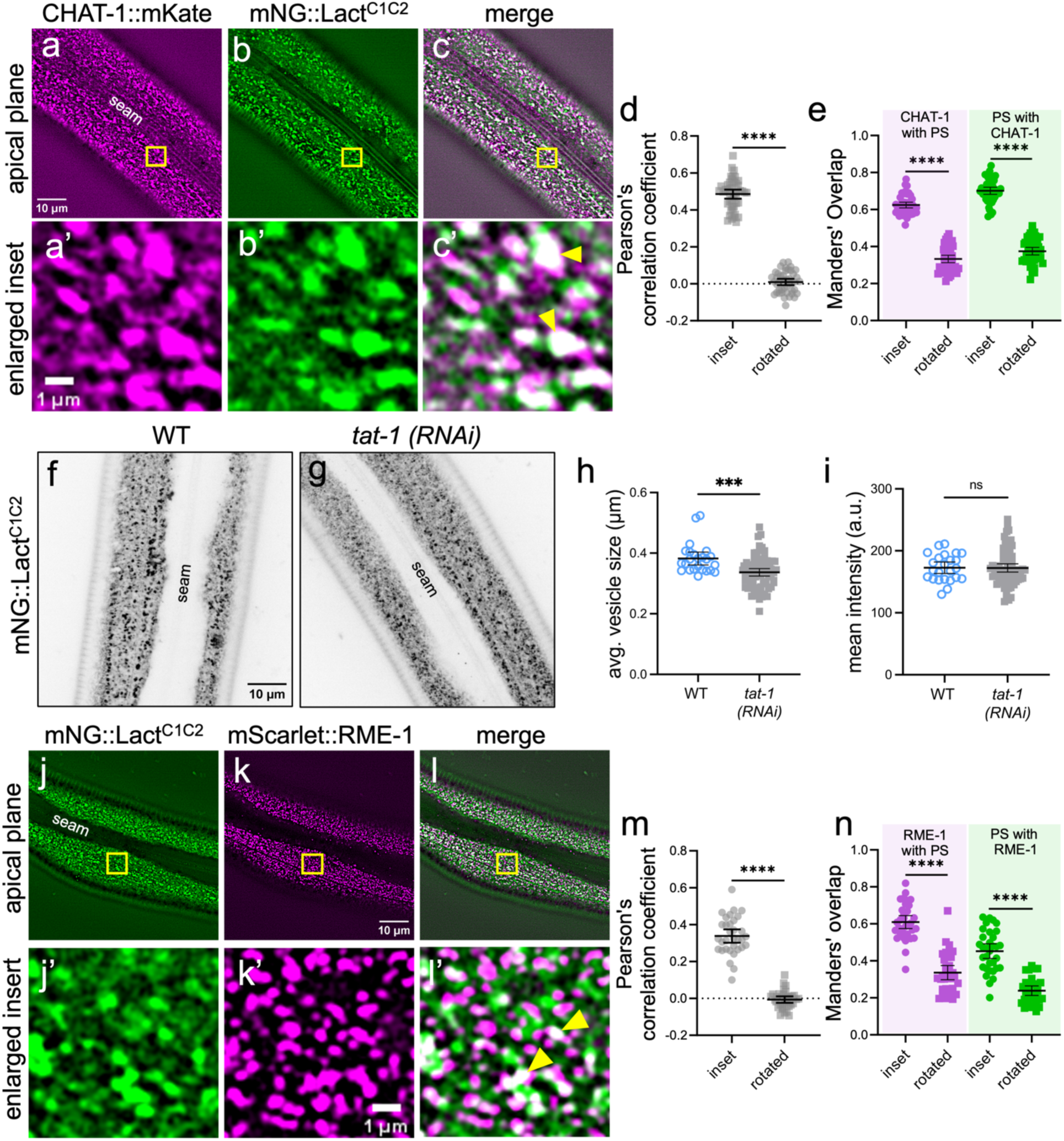
CHAT-1 and RME-1 localize to PS-labeled endosomal compartments. (a–e) Day-1 adults (n = 45) that express (a, a’) CHAT-1::mKate and (b, b’) the PS sensor P*_nekl-3_*::mNeonGreen::Lact^C1C2^ were used to determine (c, c’) colocalization, which was quantified using (d) Pearson’s correlation coefficient and (e) Manders’ overlap. (f–i) P*_nekl-3_*::mNeonGreen::Lact^C1C2^ localization in (f) wild-type versus (g) *tat-1(RNAi)* day-1 adults shows a slight reduction in (h) the average area of vesicles and no change in (i) mean intensity. (j–n) Day-1 adults (n = 33) that express (j, j’) P*_nekl-3_*::mNeonGreen::Lact^C1C2^ and (k, k’) P_hyp7_::mScarlet::RME-1 were used to determine (l, l’) colocalization, which was quantified using (m) Pearson’s correlation coefficient and (n) Manders’ overlap. Yellow boxes indicate enlarged insets. Dot plots show the mean and 95% CI. Statistical significance was determined using an unpaired t-test. ****p ≤ 0.0001, ***p ≤ 0.001; ns, not significant (p > 0.05). Raw data for all panels are available in Supplementary File 1.

**Fig. 6.**
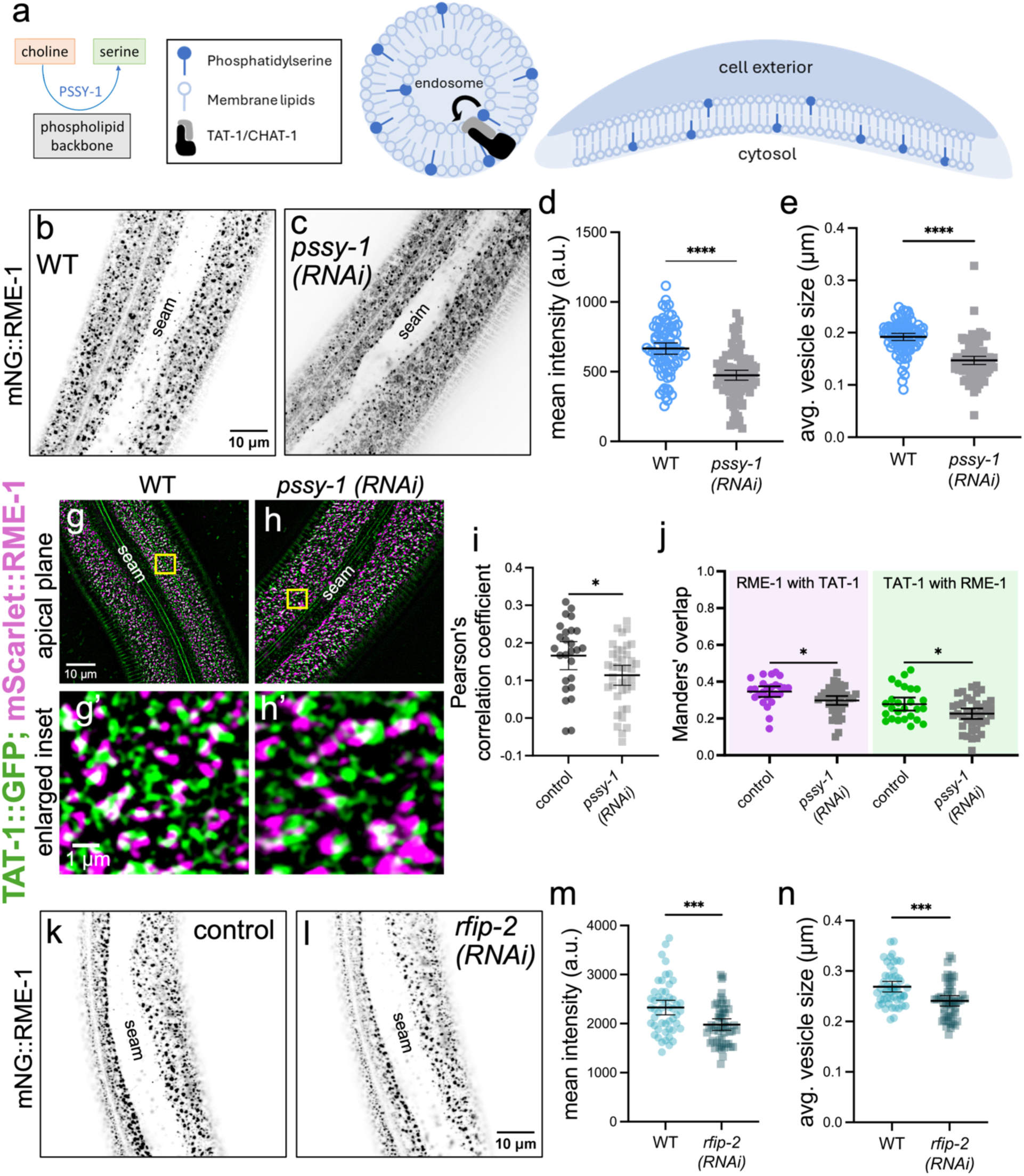
Epidermal RME-1 localization is partially dependent on PS synthase and RFIP-2. (a) Schematic showing the distribution of PS in cell membranes, the PS flippase role of TAT-1–CHAT-1, and the synthesis of PS by PSSY-1. (b–e) Representative confocal images of P_hyp7_::mNeonGreen::RME-1 in (b) WT and (c) *pssy-1(RNAi)* day-1 adults, which showed a reduced (d) mean intensity and (e) vesicle area after depletion of PS levels by *pssy-1(RNAi)*. (g–j) Colocalization of TAT-1::GFP with mScarlet::RME-1 in (g, g’) wild-type and (h, h’) *pssy-1(RNAi)* day-1 adult worms (n ≥ 26 per group). Colocalization was quantified using (i) Pearson’s correlation coefficient and (j) Manders’ overlap. Yellow boxes indicate enlarged insets. (k–n) Analysis of apical mNeonGreen::RME-1 localization in (k) WT and (l) *rfip-2(RNAi)* day-1 adults and effects on (m) mean intensity and (n) vesicle area. Dot plots show the mean and 95% CI. Statistical significance was determined using an unpaired t-test; ****p ≤ 0.0001, ***p ≤ 0.001, *p ≤ 0.05. Raw data for all panels are available in Supplementary File 1.

We next tested whether inhibition of TAT-1 affects the localization of P*_nekl-3_*::mNeonGreen::Lact^C1C2^ by knocking down TAT-1 using RNAi. We observed a modest but significant decrease in the area of puncta labeled by the sensor, consistent with TAT-1 playing a positive role in the translocation of PS to the cytoplasmic leaflet (Fig. 5f–h). In contrast, no change was observed in the total mean intensity of the biosensor (Fig. 5i), suggesting that the total levels of the sensor hadn’t changed but that its expression was more diffuse. We note that inhibition of TAT-1 would be expected to reduce, but not eliminate, PS from the cytoplasmic leaflet of endosomal membranes, consistent with our findings. Altogether, our data suggest that TAT-1 promotes the cytoplasmic translocation of PS on recycling endosomes, leading to the recruitment of RME-1.

### RME-1 localization is dependent on PS synthesis and the PS-binding protein RFIP-2

Given that inhibition of TAT-1 leads to the mis-localization of RME-1, we hypothesized that a reduction in total PS levels should also reduce RME-1 localization within apical endosomes. To test this, we carried out RNAi to inhibit PSSY-1 (<u>P</u>td<u>S</u>er <u>sy</u>nthase 1), a phosphotransferase thought to be the main producer of PS in *C. elegans* (LEVENTIS AND GRINSTEIN 2010; NILSSON *et al*. 2011). Notably, mNeonGreen::RME-1 animals injected with *pssy-1* dsRNA showed a decrease in the average area of puncta/accumulations along with a decrease in mean intensity (Fig. 6b–e), consistent with a knockdown of epidermal PS levels leading to a reduction in RME-1 recruitment to apical recycling endosomes. Furthermore, we observed a significant reduction in the extent of co-localization between TAT-1::GFP and mScarlet::RME-1 after inhibition of *pssy-1* by RNAi (Fig. 6g–j), indicating that TAT-1 does not directly recruit RME-1 but instead promotes RME-1 localization to apical endosomes through the translocation of PS.

Notably, RME-1 does not have a predicted PS-binding domain, and a previous study failed to detect binding of purified mammalian EHD1 protein to PS in vitro (NASLAVSKY *et al*. 2007). We therefore hypothesized that an additional protein may serve as a bridge to link PS at endosomal membranes with RME-1. One compelling candidate, RFIP-2, physically binds RME-1, and the human ortholog of RFIP-2, RAB11-FIP2, binds to both EHD1 and PS and acts to promote endocytic recycling (BAETZ AND GOLDENRING 2014). Consistent with this role, knockdown of RFIP-2 using RNAi in worms expressing mNeonGreen::RME-1 led to a decrease in the mean intensity and the average area of vesicles (Fig. 6k–n), similar to results obtained after depleting TAT-1 or PSSY-1. Collectively, our findings suggest that TAT-1 promotes RME-1 association with endosomal membranes in part through RFIP-2, which may directly bind and recruit RME-1.

### Suppression of *nekl* defects by inhibition of PS-associated proteins

Given that a reduction in TAT-1 activity suppressed *nekl*-associated larval arrest and led to reduced RME-1 localization at apical endosomes, we next determined whether reducing RME-1 directly would alleviate *nekl* molting defects. Notably, *rme-1(RNAi)* led to a partial suppression of *nekl-2; nekl-3* arrest (Fig. 7a). In addition, our findings for *pssy-1(RNAi)* indicated that reducing PS synthesis altered RME-1 localization in a manner similar to that of *tat-1* mutants, as did inhibition of the candidate PS–RME-1 linker protein, RFIP-2. Consistent with the above, both *pssy-1(RNAi)* and *rfip-2(RNAi)* led to significant suppression of *nekl-2; nekl-3* arrest, although, like *rme-1(RNAi)*, levels of suppression were weaker than those observed for inhibition of TAT-1 or CHAT-1 (Fig. 7a; compare with Fig. 1f, g). Collectively, these findings indicate that reduced levels of PS and RME-1 on apical endosomes likely contribute to *tat-1*–mediated suppression of *nekl* molting defects but may not fully account for the ability of *tat-1* mutants to suppress *nekl* defects.

**Fig. 7.**
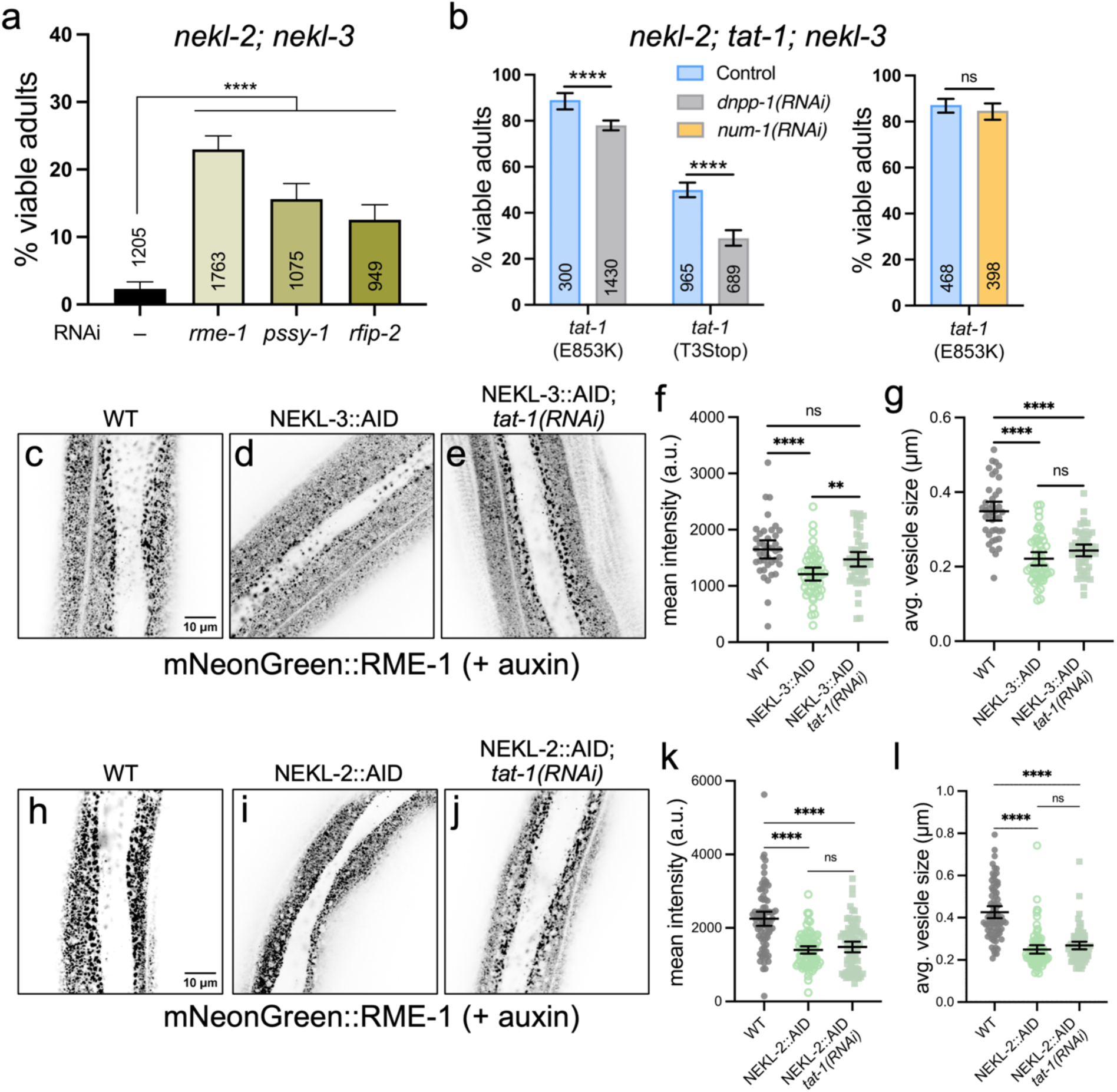
Suppression of *nekl* defects by TAT-1–RME-1 pathway components. (a) Bar plot showing the proportion of viable progeny produced by *nekl-2(fd81); nekl-3(gk894345)* animals injected with the indicated dsRNAs. (b) Bar plots showing the proportion viable adults in the progeny of *nekl-2(fd81); tat-1; nekl-3(gk894345)* hermaphrodites injected with dsRNAs targeting *dnpp-1* or *num-1a*. Bar plots show the mean and 95% CI. Statistical significance was determined by Fischer’s exact test. (c–l) Representative confocal images and analysis of auxin-treated (auxin +) 2-day-old adults expressing P_hyp7_::mNeonGreen::RME-1 in the (c, h) absence and presence of (d, e) NEKL-3::AID or (j, k) NEKL-2::AID alleles. (e, j) Animals were also treated with RNAi to knock down *tat-1*. (f, g, k, l) These effects were analyzed with respect to (f, k) mean intensity and (g, l) the average area of vesicles. NEKL-3::AID; *tat-1(RNAi)* animals showed a partial restoration of RME-1 localization as measured by mean intensity relative to NEKL-3::AID, as well as a weak trend with respect to vesicle area (p = 0.06). NEKL-2::AID and NEKL-2::AID; *tat-1(RNAi)* animals showed no significant differences. Dot plots show the mean and 95% CI (n ≥ 40 for all tests). Statistical significance was determined by unpaired t-test; ****p ≤ 0.0001, **p ≤ 0.01; ns, not significant (p > 0.05). Raw data for all panels are available in Supplementary File 1.

A previous study found that loss-of-function mutations in *dnpp-1*, an aspartyl-amino peptidase, lead to the suppression of *tat-1* mutant endocytic defects in the intestine through an unknown mechanism (LI *et al*. 2013). We reasoned that RNAi knockdown of *dnpp-1* might therefore revert the suppression of *nekl-2; tat-1; nekl-3* strains. Consistent with this, *dnpp-1(RNAi)* reduced suppression in two *nekl-2; tat-1; nekl-3* strains but did not induce larval arrest in wild type (<1%, n = 1512; Fig. 7b), consistent with DNPP-1 acting in a manner that is antagonistic to TAT-1 in the epidermis. The *C. elegans* ortholog of *Drosophila* Numb, NUM-1, has also been proposed to antagonize TAT-1 and PSSY-1 in the intestine (NILSSON *et al*. 2011). However, no reduction was observed in the suppression of *nekl-2; tat-1(E853K); nekl-3* strains when strains were treated with *num-1(RNAi)* (Fig. 7b), suggesting that NUM-1 does not antagonize TAT-1 functions in the epidermis.

In our previous studies we showed that depletion of NEKL-2 or NEKL-3 using the AID system leads to defects in multiple endocytic compartments of hyp7 in young adults (JOSEPH *et al*. 2020; JOSEPH *et al*. 2022; JOSEPH *et al*. 2023). For example, depletion of NEKL-2::AID leads to an increase in the mean intensity of RAB-5–marked early endosomes and to a decrease in the roundness of RAB-5 vesicles, whereas depletion of NEKL-3::AID leads to an increase in the mean intensity of RAB-7–marked late endosomes and a reduction in the area of RAB-7 vesicles (JOSEPH *et al*. 2023).

Moreover, depletion of NEKL-3::AID results in the abnormal accumulation of apical clathrin puncta (GFP::CHC-1) and altered localization of the clathrin-dependent cargo LRP-1 (JOSEPH *et al*. 2020). In addition, depletion of NEKL-3 leads to a decrease in the mean intensity and average area of vesicles marked with TGN-38 (JOSEPH *et al*. 2023), a resident Golgi protein that cycles between the trans Golgi, plasma membrane, and early endosomes (HUMPHREY *et al*. 1993; REAVES *et al*. 1993; ROQUEMORE AND BANTING 1998; JOSEPH *et al*. 2023). Given that loss of *tat-1* strongly suppressed *nekl* molting defects, we determined whether inhibition of TAT-1 might suppress a subset of these characterized *nekl*-associated trafficking defects. However, despite NEKL::AID depletion leading to effects that were consistent with our previous findings, we failed to observe any correction of membrane trafficking defects by *tat-1* loss of function for the above-mentioned markers (Fig. S6).

Nevertheless, given the close connection established between TAT-1 and RME-1– marked recycling endosomes in hyp7, we next looked for the effects of NEKL-2::AID and NEKL-3::AID depletion on the localization of mNeonGreen::RME-1. Notably, we found that loss of NEKL-2 or NEKL-3 led to a strong decrease in the mean intensity of RME-1 and in the area of RME-1–marked vesicles (Fig. 7c–l). Moreover, performing *tat-1(RNAi)* in NEKL-3::AID–depleted worms led to a substantial increase in the mean intensity of mNeonGreen::RME-1 and to a marginally significant increase (p = 0.06) in the average area of RME-1–marked vesicles relative to the NEKL-3::AID–depleted worms with functional *tat-1* [(Fig. 7d–g). In contrast, *tat-1(RNAi)* in NEKL-2::AID showed at most a weak trend toward restoring mNeonGreen::RME-1 localization, which did not reach statistical significance (Fig. 7i–l). Collectively, our data suggest that *tat-1* mutant–mediated suppression of *nekl* molting and trafficking defects is likely to occur at least in part through its role in the regulation of RME-1.

## DISCUSSION

Here we report the identification of mutations in *tat-1* as suppressors of *nekl*-associated molting defects. Previous work in *C. elegans* and mammals has shown that TAT-1/ATP8A2, a PS flippase, and its chaperone, CHAT-1/CDC50, regulate endocytosis by maintaining PS asymmetry on the outer leaflet of endosomes and the plasma membrane, thereby promoting processes including membrane tubulation and vesicle scission (DARLAND-RANSOM *et al*. 2008; RUAUD *et al*. 2009; CHEN *et al*. 2010; BEST *et al*. 2019). Consistent with the above, we found that inhibition of *chat-1* strongly suppressed *nekl* defects and observed strong colocalization of endogenously tagged TAT-1 and CHAT in the epidermis. Previous work on *C. elegans* TAT-1 and CHAT-1 focused largely on their functions in germline cell engulfment and endocytic trafficking in intestinal cells (ZULLIG *et al*. 2007; DARLAND-RANSOM *et al*. 2008; RUAUD *et al*. 2009; CHEN *et al*. 2010; MAPES *et al*. 2012). Because loss of TAT-1 and CHAT-1 in the intestine lead to the formation of enlarged hybrid endosomal compartments, it was proposed that TAT-1 and CHAT-1 may promote the maturation of late endosomes and lysosomes (RUAUD *et al*. 2009). Although we observed an increase in the intensity and area of puncta corresponding to early (RAB-5), late (RAB-7), and recycling (RAB-10 and RAB-11) endosomes, we failed to detect enlarged hybrid compartments in the epidermis of *tat-1* mutants. We also note that in contrast to previous studies that used multi-copy TAT-1 and CHAT-1 reporters driven by intestinal-specific promoters (RUAUD *et al*. 2009; CHEN *et al*. 2010), we detected minimal TAT-1 and CHAT-1 expression in intestinal cells in our CRISPR-tagged strains. Collectively, these results suggest that TAT-1, along its impact on PS distribution, likely has distinct functions within different cell types.

In contrast to the increased expression of RAB markers in *tat-1* mutants, we observed a substantial decrease in the mean intensity and area of RME-1–marked compartments in *tat-1* mutants. Consistent with this, we detected strong colocalization of TAT-1 with RME-1 in mesh-like structures and with RAB-5–positive early endosomes in round, punctate structures in the apical portion of hyp7. These results suggest that the primary action of TAT-1–CHAT-1 in the epidermis takes place at or near the transition step between early and recycling endosomes, although low or undetectable levels of activity could be present elsewhere. Our findings are analogous to a previous study in *C. elegans* intestinal cells in which TAT-1 was found to colocalize with RME-1 in apical regions and in which loss of *tat-1* led to a decrease in apical RME-1 localization (CHEN *et al*. 2010). However, whereas loss of *tat-1* and *chat-1* also leads to a strong reduction in RME-1 at basolateral membranes in intestinal cells (RUAUD *et al*. 2009; CHEN *et al*. 2010; LI *et al*. 2013), we failed to detect RME-1, TAT-1, or CHAT-1 at basolateral membranes in the epidermis. These findings from *C. elegans* are consistent with knockdown of ATP8A1/2 in mammalian cell culture, which causes a reduction in the localization of EHD1 at perinuclear recycling endosomes (LEE *et al*. 2015).

Our study made use of a cytoplasmic hyp7-specific PS sensor, which was localized to TAT-1– and RME-1–positive apical recycling endosomes. In addition, our co-localization studies suggested that PS may be localized to portions of early endosomes. Consistent with TAT-1 functioning as a PS flippase, *tat-1* inhibition reduced localization of the PS sensor at apical puncta without altering overall expression levels of the sensor, suggesting that the signal derived from the PS sensor had become more diffuse. Altogether, our cumulative data indicate that loss of TAT-1 flippase activity leads to the reduced recruitment of RME-1 to apical endosomes, which is likely mediated by the PS- and RME-1–binding protein RFIP-2.

One key question raised by our findings relates to the mechanism by which loss of TAT-1 suppresses *nekl* molting defects. Several suppressors identified by our genetic screen, including the F-BAR domain–containing protein FCHO-1; SID-3, an activated CDC42 protein kinase; and members of the AP2 clathrin-adapter complex, appear to suppress *nekl* molting defects by partially correcting *nekl-*associated defects in membrane trafficking (LAŽETIĆ *et al*. 2018; JOSEPH *et al*. 2020). By contrast, we have failed to identify direct roles in trafficking for several other *nekl* suppressors including the conserved ADAM protease family member ADM-2, the Na^+^/K^+^ cation pump CATP-1, and mutations that affect entry into the dauer pathway (BINTI *et al*. 2022; JOSEPH *et al*. 2022; BINTI *et al*. 2024a). Consistent with TAT-1 acting through an established trafficking pathway, we demonstrated that RME-1 localization is at least partly dependent on RFIP-2, PSSY-1, and TAT-1, and that inhibition of any of these proteins suppresses *nekl* molting defects. However, the quantitatively stronger suppression observed after inhibition of TAT-1 suggests that *tat-1*–mediated suppression may act through other proteins and lipid species.

We also found that depletion of NEKL-2::AID or NEKL-3::AID in the presence of auxin led to a decrease in the mean intensity and average area of vesicles labeled by RME-1, a phenotype similar to what we observed in *tat-1* single mutants. We were therefore surprised by our finding that *tat-1(RNAi)* partially restored RME-1 localization in NEKL-3::AID–depleted worms. Although this result does support the general conclusion that loss of TAT-1 suppresses *nekl* molting defects through a trafficking-based mechanism, further studies will be needed to understand the precise means by which this occurs.

Another question raised by our studies concerns the discrepancy between the strong genetic suppression of molting defects conferred by *tat-1* inhibition versus the relatively weak suppression of RME-1 localization defects in NEKL::AID-depleted worms. In this regard, it is worth noting that whereas *tat-1* mutations and RNAi strongly suppressed hypomorphic alleles of *nekl-2* and *nekl-3*, TAT-1 inhibition led to little or no suppression of *nekl* null mutations. Given that the auxin-based AID system leads to the near-complete degradation of tagged proteins (ZHANG *et al*. 2015; JOSEPH *et al*. 2020; ASHLEY *et al*. 2021; PHANINDHAR AND MISHRA 2023), it is perhaps not surprising that *tat-1* inhibition did not strongly suppress RME-1 localization defects in NEKL-2::AID and NEKL-3::AID strains.

In summary, our findings highlight the strength of suppressor genetic approaches for identifying regulators of endocytic trafficking. More specifically, our studies provide new links between conserved NIMA-related kinases and several known membrane trafficking proteins with links to PS regulation and enrichment including RME-1, RFIP-2, and TAT-1. Our studies also highlight differences in the roles of lipid-modifying enzymes across cell types and during development.

## DATA AVAILABILITY

Raw data for all figures are provided in Supplementary File 1. Any additional information is available upon request.

## ACKNOWLEDGEMENTS

We thank Amy Fluet for editing this manuscript and Barth Grant for providing strains. Some strains were provided by the CGC, which is funded by NIH Office of Research Infrastructure Programs (P40 OD010440).

## SUPPLEMENTARY FIGURES

## SUMMARY OF SUPPLEMENTAL FILES

**File S1. Raw data files for all figures (Excel)**

Compilation of raw data used in this study.

**File S2. Additional technical information (Excel)**

Strains: List of all the strains used in this study.

RNAi: List of RNAi primers used in this study.

CRISPR: List of all CRISPR repair templates, guide RNAs, screening primers, and restriction enzymes used in this study.

---

**Fig. S1.**
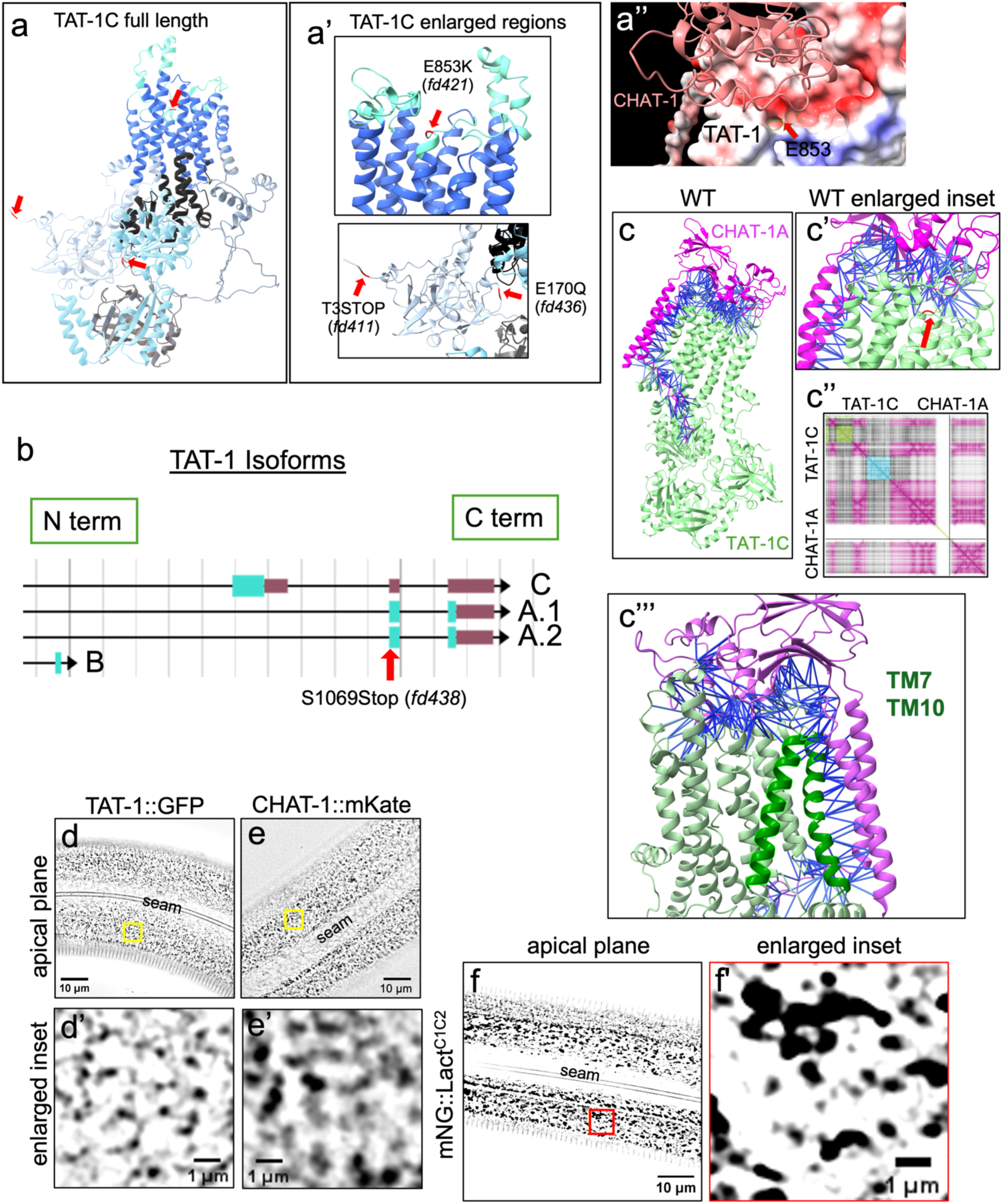
Predicted TAT-1–CHAT-1 interactions and expression of TAT-1, CHAT-1, and a PS sensor in the epidermis. (a–a’’) AlphaFold2/3 predictions of TAT-1 rendered by ChimeraX showing locations of the relevant mutations (red arrows). TAT-1 is rendered in (a’’) surface-charge format (red acidic–blue basic) with CHAT in ribbon format; E853 is indicated by the red arrow and green lines. (b) TAT-1 isoforms (C, A.1, A.2, B) as depicted on WormBase. The CRISPR-engineered Stop mutation (*fd438*) specifically affects the coding portion of the C terminus of the A isoforms. (c, c’) AlphaFold3 multimer prediction of binding between TAT-1C and CHAT-1A (interface predicted template modelling score (ipTM) = 0.88). Blue lines represent predicted contact regions (predicted aligned error (PAE) < 5 Å, backbone distance < 5 Å). Red arrow indicates location of E853. (c’’) Two-dimensional PAE diagram colored by PAE domain as rendered by ChimeraX. Magenta regions in upper right and lower left quadrants indicate regions of TAT-1 and CHAT-1 predicted to form a structurally coherent multi-protein domain. (d–e’) Representative deconvoluted confocal images of day-1 adultss showing endogenous (CRISPR-tagged) (d, d’) TAT-1::GFP and (e, e’) CHAT-1::mKate in the apical region of hyp7. Yellow boxes in (d, e) indicate the enlarged insets in (d’, e’). (f, f’) The genetically encoded PS sensor P*_nekl-3_*::mNeonGreen::Lact^C1C2^ exhibits an apical mesh-like localization pattern in the epidermis. Red box in (f) indicates the enlarged inset in (f’). Raw data for all panels are available in Supplementary File 1.

**Fig. S2.**
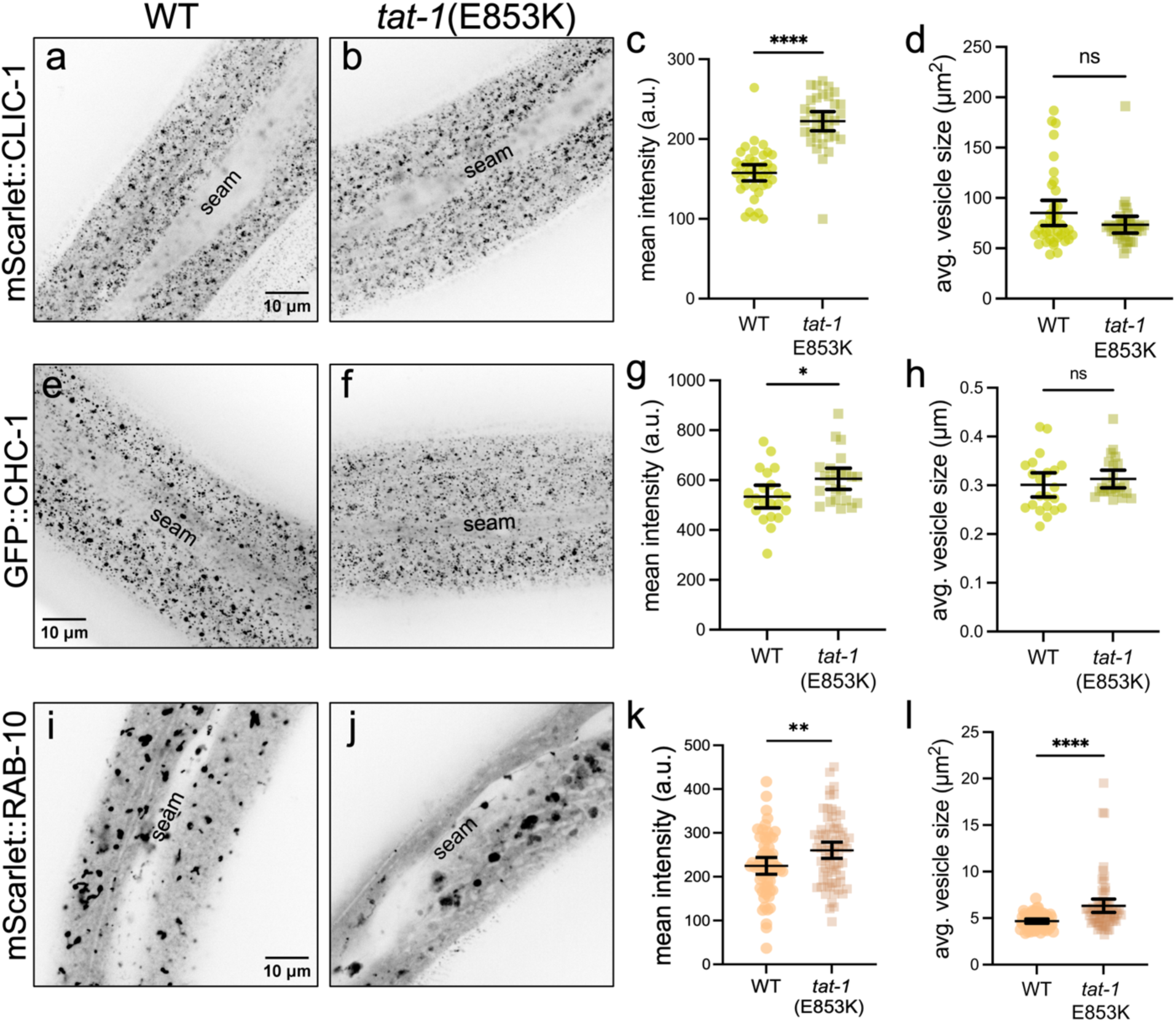
*tat-1* loss of function affects size and intensity of clathrin-coated pits and RAB-10–positive endosomes. (a, b, e, f, i, j) Representative confocal images showing day-1 adults expressing (a, b) mScarlet::CLIC-1, (e, f) GFP::CHC-1, or (i, j) P_hyp-7_::mScarlet::RAB-10 in (a, e, i) WT and (b, f, j) *tat-1* (E853K) animals. (c, d, g, h, k, l) Dot plots show the (c, g, k) mean intensity, (d) area of puncta, and (h, l) average area of vesicles. The mean and 95% CI are indicated. Statistical significance was determined using an unpaired t-test; ****p ≤ 0.0001, **p ≤ 0.01, *p ≤ 0.05; ns, not significant (p > 0.05). The number of animals quantified in each group is indicated. Raw data for all panels are available in Supplementary File 1.

**Fig. S3.**
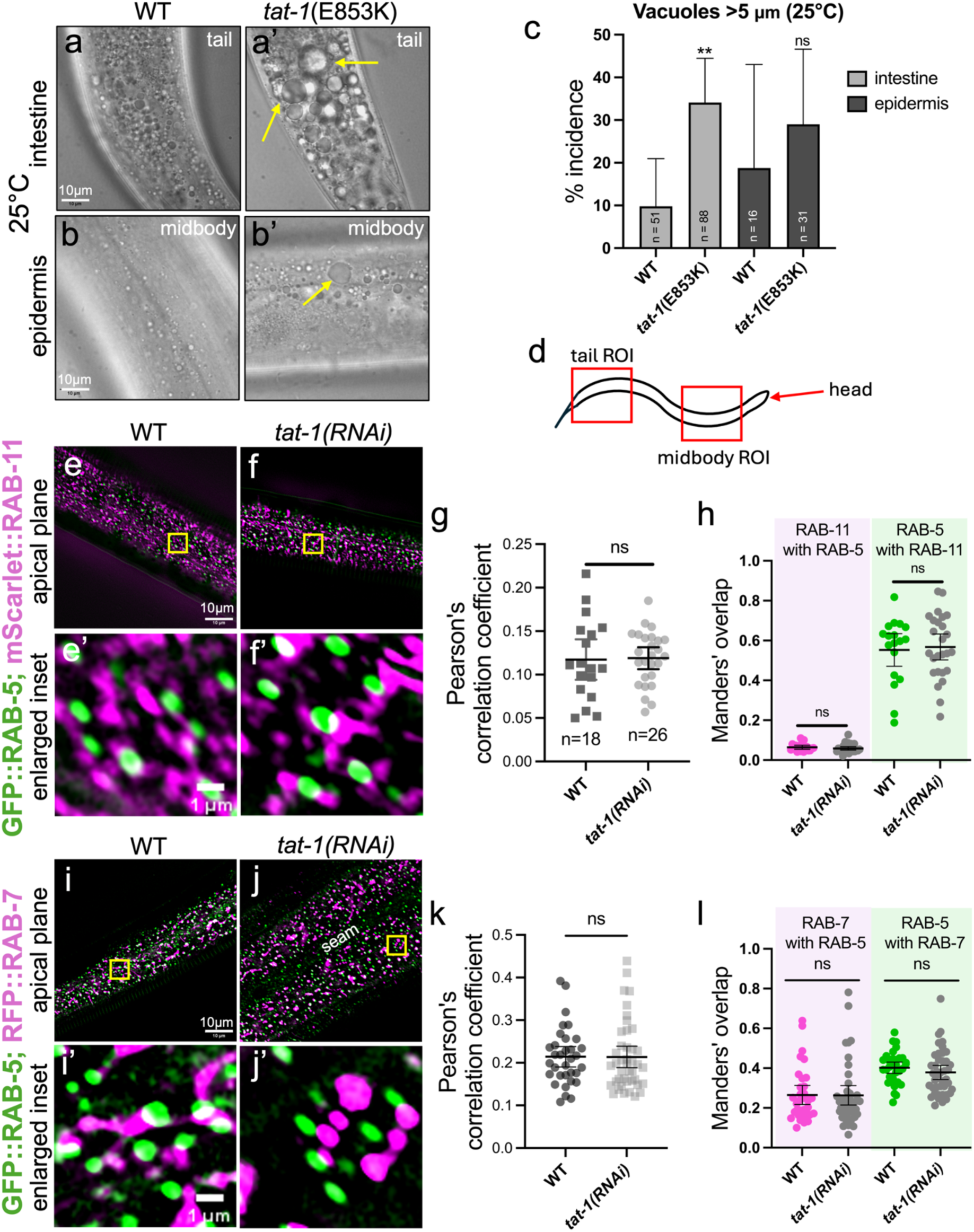
Assessment of *tat-1*–mediated vacuolation and RAB-5–RAB-11 and RAB- 5–RAB-7 hybrid endosome phenotypes. (a–b’) Representative bright-field images of (a, a’) intestine and (b, b’) epidermis in (a, b) wild-type and (a’, b’) *tat-1*(E853K) day-1 adults. Yellow arrows indicate vacuoles in the intestine and epidermis of *tat-1*(E853K) animals. (c) Quantification of vacuole phenotype at 25°C in the intestine and epidermis of wild-type and *tat-1*(E853K) animals. (d) Schematic shows regions of interest (ROIs; red boxes) of *C. elegans* that were imaged for (a–c). (e–f’, i–j’) Co-expression of (e–f’) RAB-5–RAB-11 and (i–j’) RAB-5–RAB-7 in wild-type control and *tat-1(RNAi)* day-1 adults. Yellow boxes indicate enlarged insets. (g, h, k, l) Colocalization was quantified using (g, k) Pearson’s correlation coefficient and (h, l) Manders’ overlap. Dot plots show the mean and 95% CI. Statistical significance was determined using an unpaired t-test; ns, not significant (p > 0.05). Raw data for all panels are available in Supplementary File 1.

**Fig. S4.**
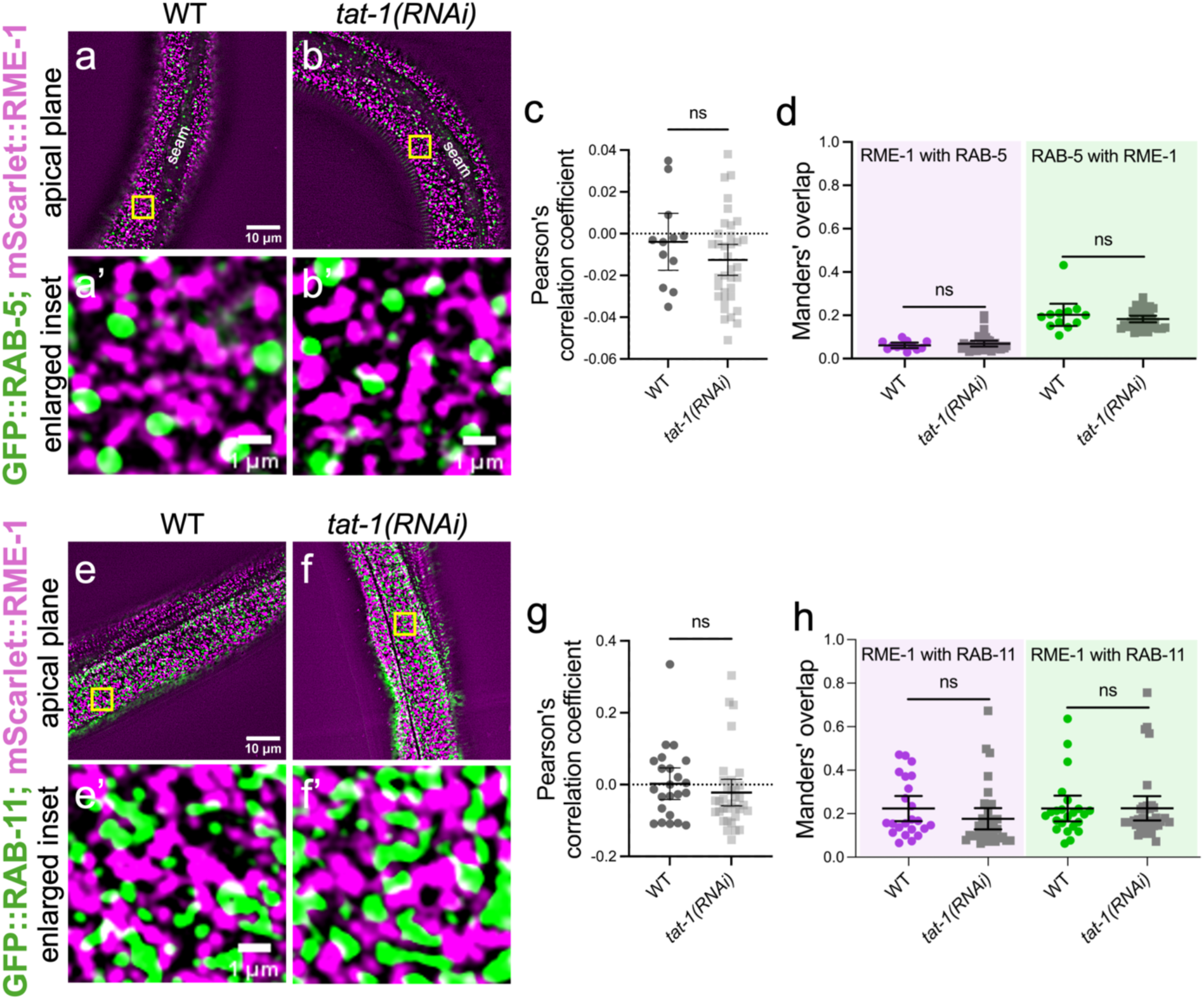
Assessment of *tat-1*–mediated RAB-5–RME-1 and RAB-11–RME-1 hybrid-endosome phenotypes. (a–b’, e–f’) Co-expression of (a–b’) RAB-5–RME-1 and (e–f’) RAB-11–RME-1 in wild-type control and *tat-1(RNAi)* day-1 adults. Yellow boxes indicate enlarged insets. (c, d, g, h) Colocalization was quantified using (c, g) Pearson’s correlation coefficient and (d, h) Manders’ overlap. Dot plots show the mean and 95% CI. Statistical significance was determined using an unpaired t-test; ns, not significant (p > 0.05). Raw data for all panels are available in Supplementary File 1.

**Fig. S5.**
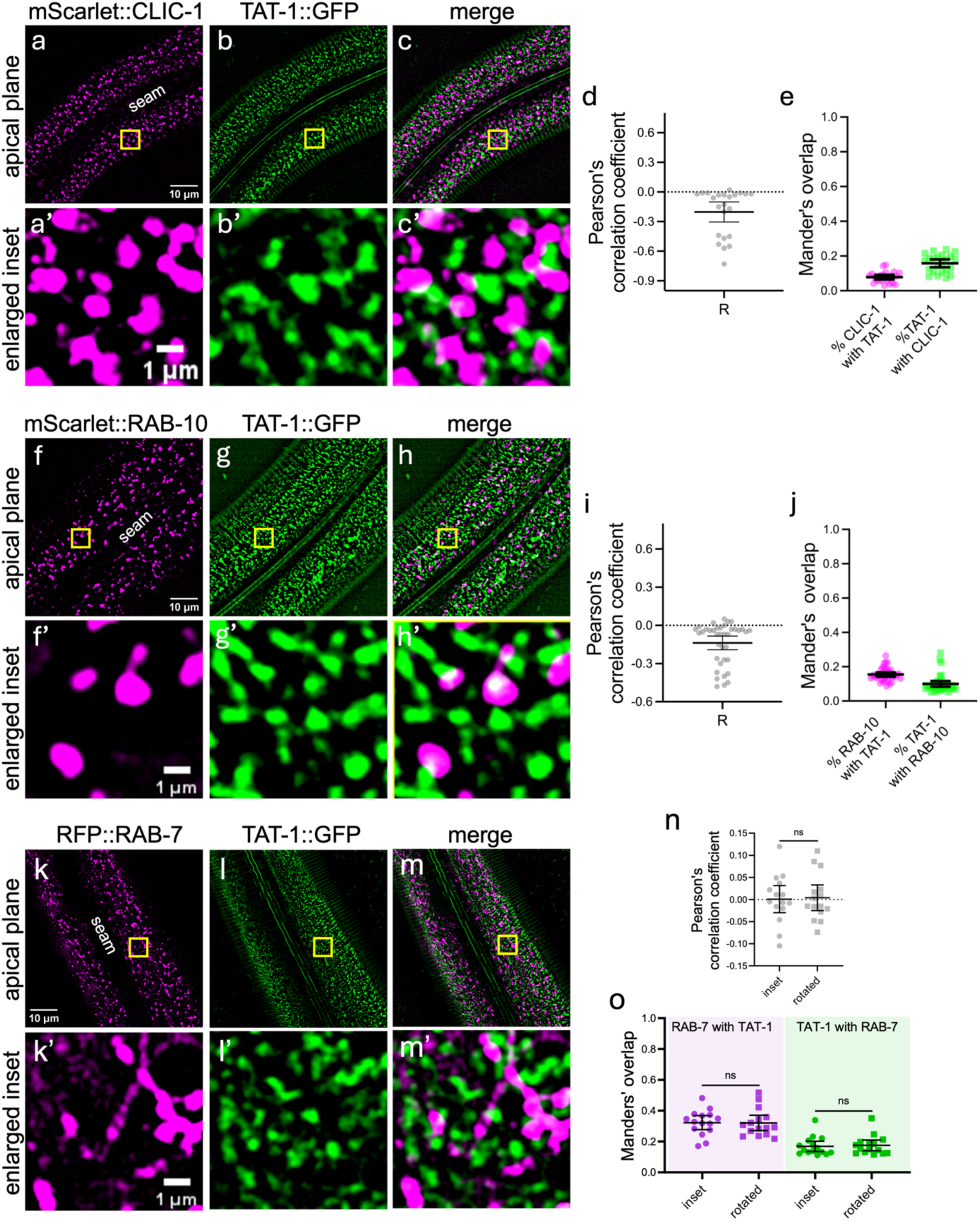
TAT-1 colocalization with additional endocytic markers. (a–c’, f–h’, k–m’) Representative confocal images of TAT-1::GFP co-expressed with (a–c’) mScarlet::CLIC-1 (n = 23), (f–h’) P_hyp7_::mScarlet::RAB-10 (n = 38), and (k–m’) RFP::RAB-7 (n = 15) in day-1 adults. Yellow boxes indicate enlarged insets. Colocalization was quantified using (d, i, n) Pearson’s correlation coefficient and (e, j, o) Manders’ overlap. Dot plots show the mean and 95% CI. Statistical significance in (n, o) was determined using an unpaired t-test; ns, not significant (p > 0.05). Raw data for all panels are available in Supplementary File 1.

**Fig. S6.**
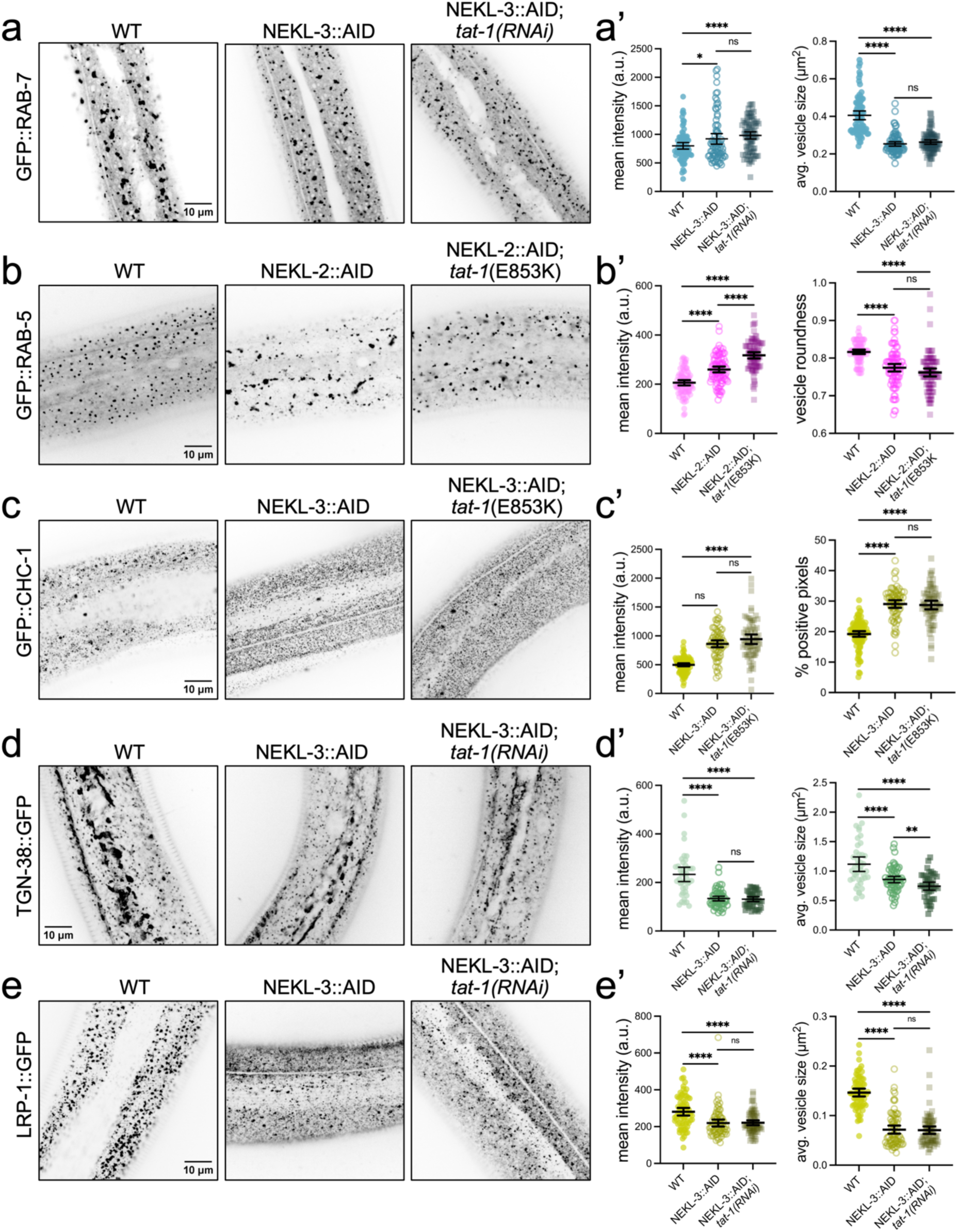
Inhibition of *tat-1* does not suppress several trafficking defects associated with the depletion of NEKL-2 or NEKL-3. (a–e) Representative confocal images of day-2 adults s that express markers associated with endocytic compartments after auxin treatment to deplete NEKL-3 or NEKL-2 and in the presence and absence of *tat-1(RNAi)*. Animals expressed (a) P_hyp7_::RFP::RAB-7, (b) P*_rab-5_*::GFP::RAB-5, (c) GFP::CHC-1, (d) P_hyp7_::TGN-38::GFP, and (e) LRP-1::GFP. (a’–e’) Animals were analyzed for mean intensity and (a’, d’, e’) average area of vesicles, (b’) vesicle roundness, or (c’) percent positive pixels. Dot plots show the mean and 95% CI. Statistical significance was determined by unpaired t-test; ****p ≤ 0.0001, **p ≤ 0.01, *p ≤ 0.05; ns, not significant (p > 0.05). Raw data for all panels are available in Supplementary File 1.

